# The YTHDF proteins shape the brain gene signatures of Alzheimer’s disease

**DOI:** 10.1101/2024.10.23.619425

**Authors:** Shinya Tasaki, Denis R. Avey, Nicola A. Kearns, Chunjiang Yu, Sashini De Tissera, Himanshu Vyas, Lin Cheng, Jishu Xu, Artemis Iatrou, Daniel J. Flood, Wenlong Li, Lisa L. Barnes, Katherine Rothamel, Aliza P Wingo, Thomas S Wingo, Nicholas T. Seyfried, Chuan He, Philip L. De Jager, Gene Yeo, Chris Gaiteri, David A. Bennett, Yanling Wang

## Abstract

The gene signatures of Alzheimer’s Disease (AD) brains reflect an output of a complex interplay of genetic, epigenetic, epi-transcriptomic, and post-transcriptional regulation., yet the dominant factor shaping these signatures remains unclear. To identify the most significant factor that shapes the AD brain signatures, we integrated cellular and molecular features with differential gene expression in in an explainable machine learning framework. Our result indicates that YTHDF proteins, the canonical readers of N6-methyladenosine RNA modification (m6A), are the most influential predictors of the AD brain signatures. We then show that protein modules containing YTHDFs are downregulated in human AD brains, and knocking down and pharmacologically inhibiting YTHDFs in iPSC-derived 2D and 3D neuronal models recapitulate the AD-associated transcriptional signatures. Furthermore, eCLIP-seq analysis revealed that YTHDF proteins influence AD signatures through both m6A-dependent and independent pathways. These results highlight the central role of YTHDF proteins in shaping the gene signatures of AD brains.

## Main

Many RNA expression changes have been identified in Alzheimer’s disease (AD) brains^1–3^. These changes could be influenced by multiple factors, such as genetic, epigenetic, epi-transcriptomic, and post-transcriptional mechanisms. These regulatory factors often interact in a cell-type-specific manner, contributing to the complexity and heterogeneity of the disease process^4^. However, it remains unclear which of these factors is primarily responsible for driving the AD gene signature and, ultimately, how to halt the progression of molecular dysregulation in AD. Existing purely data-driven approaches, such as co-expression networks^1–3^, are valuable but can be difficult to link to specific genomic mechanisms. In contrast, knowledge-based approaches, such as genomic interaction databases (e.g., transcription factor-DNA, RNA-RNA, protein-RNA interactions) generated by consortia^5–7^ hold valuable insights; however, their integration with gene expression data has often been insufficient for defining overarching disease mechanisms. These challenges have recently motivated the application of explainable machine learning approaches^8^ to efficiently integrate multi-omics data, identify expression patterns, and predict top regulators and disease mechanisms.

In this study, we integrate these two approaches using an explainable machine learning model to discover regulatory features of gene signatures associated with cognitive decline in AD. Our analysis predicts that YTH domain-containing family proteins (YTHDFs), the classical reader proteins of N6-methyladenosine (m6A) RNA modification, shape the gene signatures of AD brains. We then validate our predictions using global proteomics analysis and perturbation experiments. To examine whether YTHDF proteins act through their canonical pathway of recognizing m6A sites, we conducted enhanced crosslinking and immunoprecipitation followed by sequencing (eCLIP-seq) to simultaneously examine YTHDF binding sites and m6A modification sites. Our results suggest a combined function of YTHDF proteins in shaping AD gene signatures via either m6A-dependent or m6A-independent pathways, offering potential therapeutic targets that specifically address molecular changes in AD brains.

## Results

### m6A readers are the most influential predictors for differentially expressed genes in AD

Transcriptome analysis of bulk tissue from large-scale brain cohorts has identified many differentially expressed genes (DEGs) associated with AD phenotypes, such as neuropathologies and cognitive decline^1^ (**Fig. 1a,b**). Since these gene signatures potentially reflect the integral output of multifaceted regulatory mechanisms, it remains challenging to untangle this complexity, uncover the most influential mechanisms shaping these specific gene signatures, and identify their cell types of action.

**Figure 1.**
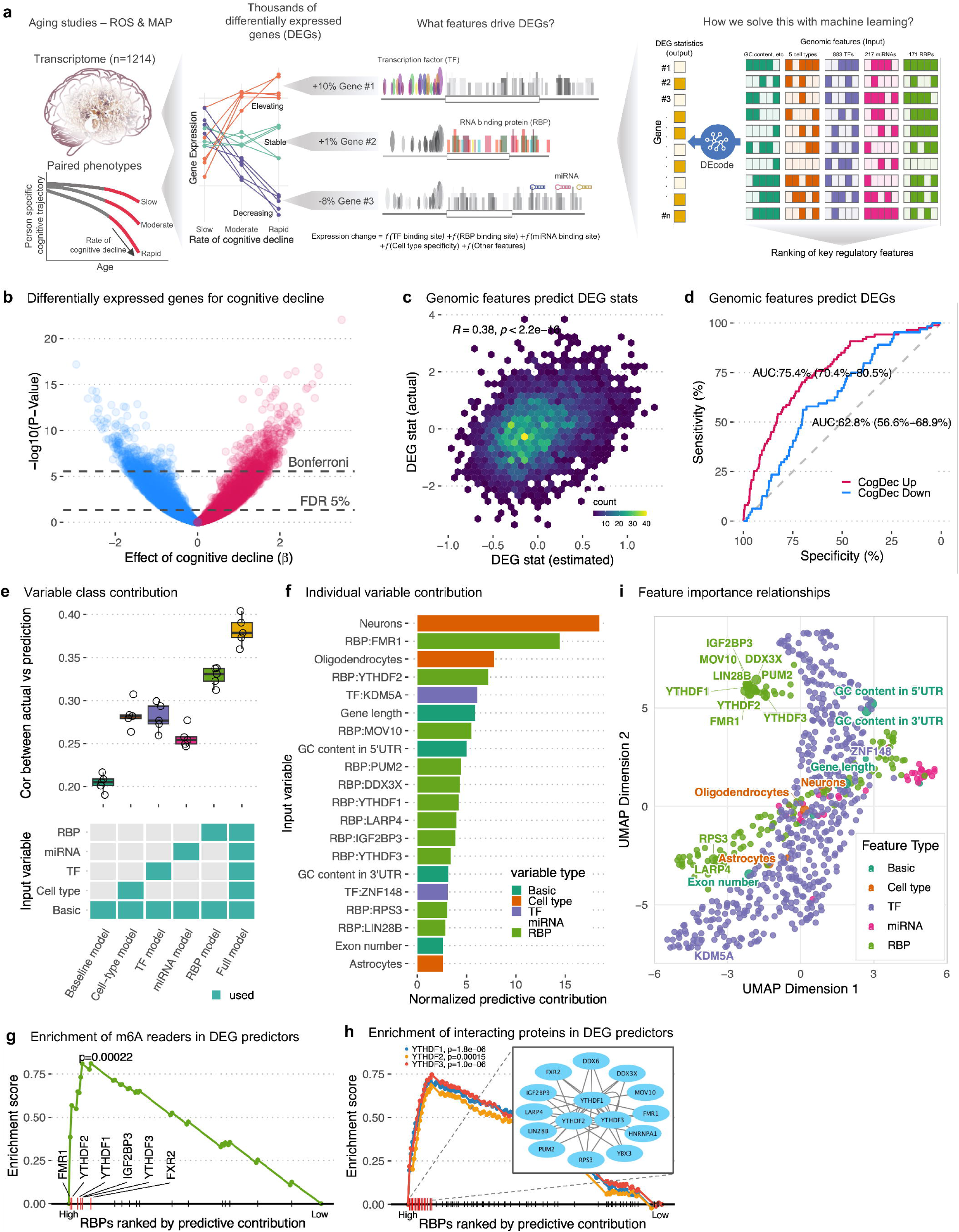
Regulatory determinants of differentially expressed genes for cognitive decline in DLPFC

To address this challenge, we built a machine learning framework, DEcode-tree, that explains DEGs related to the rate of cognitive decline in AD. With interpretability as our foundation, we compiled biophysical features of basic gene information (number of exons, 5’ UTR length, transcript length, GC content) and regulatory features of binding sites (833 transcription factors, 172 RNA-binding proteins, and 217 miRNAs), along with gene expression profiles of major neural cell types (neurons, astrocytes, microglia, oligodendrocytes, and oligodendrocyte precursor cells). Through the integration of these diverse data types, the model captures the multifaceted regulatory landscape influencing gene expression changes associated with cognitive decline. Crucially, the model predicts differential-expression status for genes never seen during training using a gene-holdout paradigm, enabling discovery of transferable regulatory principles rather than memorizing gene-specific patterns. We applied DEcode-tree to the DLPFC RNA-seq data from 761 ROSMAP participants (**Supplementary Table 1**). By contrasting gene expression levels with the rate of cognitive decline, we identified 6,855 DEGs at an FDR of 5%, and 775 DEGs after Bonferroni correction (**Fig. 1b**, and **Supplementary Table 2**). Our model predicted DEGs with an overall correlation of R=0.38 and P=6.66×10^−120^ to the actual DEG statistics (**Fig. 1c**). The AUC (Area Under the Curve) value was 0.75 (90% CI: 0.70-0.81) for upregulated genes and 0.63 (90% CI: 0.57-0.69) for downregulated genes (**Fig. 1d**). The model achieved a 47% improvement in predictive performance over our previous method^8^ (**Supplementary Fig. 1a**), reaching an overall correlation of R=0.38. This represents approximately 60% of the empirical maximum observed when comparing t-statistics between independent batches of expression data (R∼0.63, **Supplementary Fig. 1b**). This empirical ceiling reflects the inherent biological variability and technical noise in gene expression measurements, positioning our model’s performance as robust within the constraints of gene regulatory complexity.

To identify the most impactful features in our model, we conducted an ablation analysis by systematically removing each input, retraining the model, and then evaluating the model’s performance. Notably, the model trained with RNA-binding protein (RBP) binding sites performed better (R=0.33, median absolute deviation (MAD) = 0.01) than models trained with other inputs such as transcription factor (TF) binding sites (R=0.28, MAD = 0.03), in predicting the cognitive decline gene signature (**Fig. 1e**), highlighting the importance of post-transcriptional regulation via RBPs in AD pathophysiology.

To narrow down the essential individual features in our model, we computed the SHAP score^9^, a value that represents the contribution of each feature to the prediction. We found that the most predictive cell types were neurons, oligodendrocytes, and astrocytes (**Fig. 1f**). Aligning with our ablation analysis, the RBPs are the essential predictors (**Fig. 1f** and **Supplementary Table 3**), and among the RBPs, the canonical reader proteins for m6A modification—including FMR1 and YTHDF1, YTHDF2, and YTHDF3—were most influential predictors (**Fig. 1f**). These RBPs are preferentially bound to up-regulated differentially expressed genes, with approximately 75% of upregulated genes bound by YTHDF1 and YTHDF2, and slightly lower but significant binding percentages observed for YTHDF3 and FMR1 (**Supplementary Fig. 1c**). Because m6A is one of the most common RNA modifications and plays a crucial role in regulating RNA stability, translation, and degradation, our finding prompted us to examine the performance of all m6A reader proteins in our model. To do that, we conducted gene-set enrichment analysis using 23 known m6A readers^10^ and detected their significant enrichment among all RBP predictors (P=2.2 × 10^−4^) (**Fig. 1g**). Given that YTHDFs often interact with other proteins to regulate RNA metabolism^11^, we then calculated the enrichment of their interacting proteins among the RBP predictors. We found that the interacting proteins of YTHDF1 (P=1.8 × 10^−6^), YTHDF2 (P=1.5 × 10^−4^), and YTHDF3 (P=1.0 × 10^−6^) were also enriched (**Fig. 1h**). To further understand the relationships between predictors in our model, we analyzed pairwise SHAP correlations using UMAP visualization (**Fig. 1i**). This analysis revealed that m6A-related RBPs form a distinct, tightly clustered group, confirming their coordinated contribution to differential expression and supporting their known protein-protein interactions. These findings further underscore the importance of m6A-related RBPs in regulating gene expression related to cognitive decline in AD.

To further confirm the impact of RBPs on the model’s performance in predicting DEGs associated with cognitive decline, we trained the model on an independent batch of transcriptome data (n□=□453). We achieved highly congruent results (**Supplementary Fig. 1d**). Furthermore, we applied the DEcode-tree to predict DEGs associated with AD diagnosis (**Supplementary Table 2**) and observed a similarly strong predictive power of RBPs and m6A reader proteins (**Supplementary Fig. 1e**). Collectively, our findings suggest that post-transcriptional regulation via RBPs, especially m6A readers, shapes AD molecular phenotypes.

### YTHDF-containing protein modules explain most differentially expressed genes in AD

Based on the observation that RNA targets of m6A reader RBPs are differentially expressed in AD, we next examined the dysregulation of the RBPs themselves. Since RBP transcripts must be translated into proteins, to subsequently bind RNA, we analyzed the protein levels of m6A-related RBPs in AD DLPFC using global proteomics data from 596 ROSMAP individuals (549 of which had paired transcriptome data) (**Supplementary Table 4**). As RBPs often interact with other proteins to mediate various post-transcriptional processes, we first constructed protein co-abundance modules from 7,640 detected global proteins (**Supplementary Table 5**), and then associated the average expression of each module with rapid cognitive decline and AD clinical diagnosis (**Fig. 2a**). Among the 33 protein modules (range: 35 to 567 proteins; median: 194), 16 modules were significantly associated with cognitive decline, 16 modules were associated with AD diagnosis (FDR at 5%), and 15 modules were associated with both traits (**Fig. 2b** and **Supplementary Table 6**). To provide broader molecular context for RBPs, we examined which modules contained m6A-related proteins and the key DEG predictors identified by DEcode-tree (**Fig. 1f** and **Supplementary Table 3**). m6A writers were enriched in module m3 (463 proteins, P=6.0 × 10^−5^), but this module was not associated with either trait. Notably, we found that DEG predictors (P=1.3 × 10^−6^) and cytoplasmic m6A readers (P=6.2 × 10^−4^) were both significantly enriched in m58 (125 proteins) (**Fig. 2c** and **Supplementary Table 6**), and associated with cognitive decline and AD diagnosis. Moreover, the sets of interacted proteins of YTHDF1 (P=6.5 × 10^−11^), YTHDF2 (P=5.0 × 10^−14^), and YTHDF3 (P=1.2 × 10^−7^) were all enriched in m58 (**Supplementary Table 6**).

**Figure 2.**
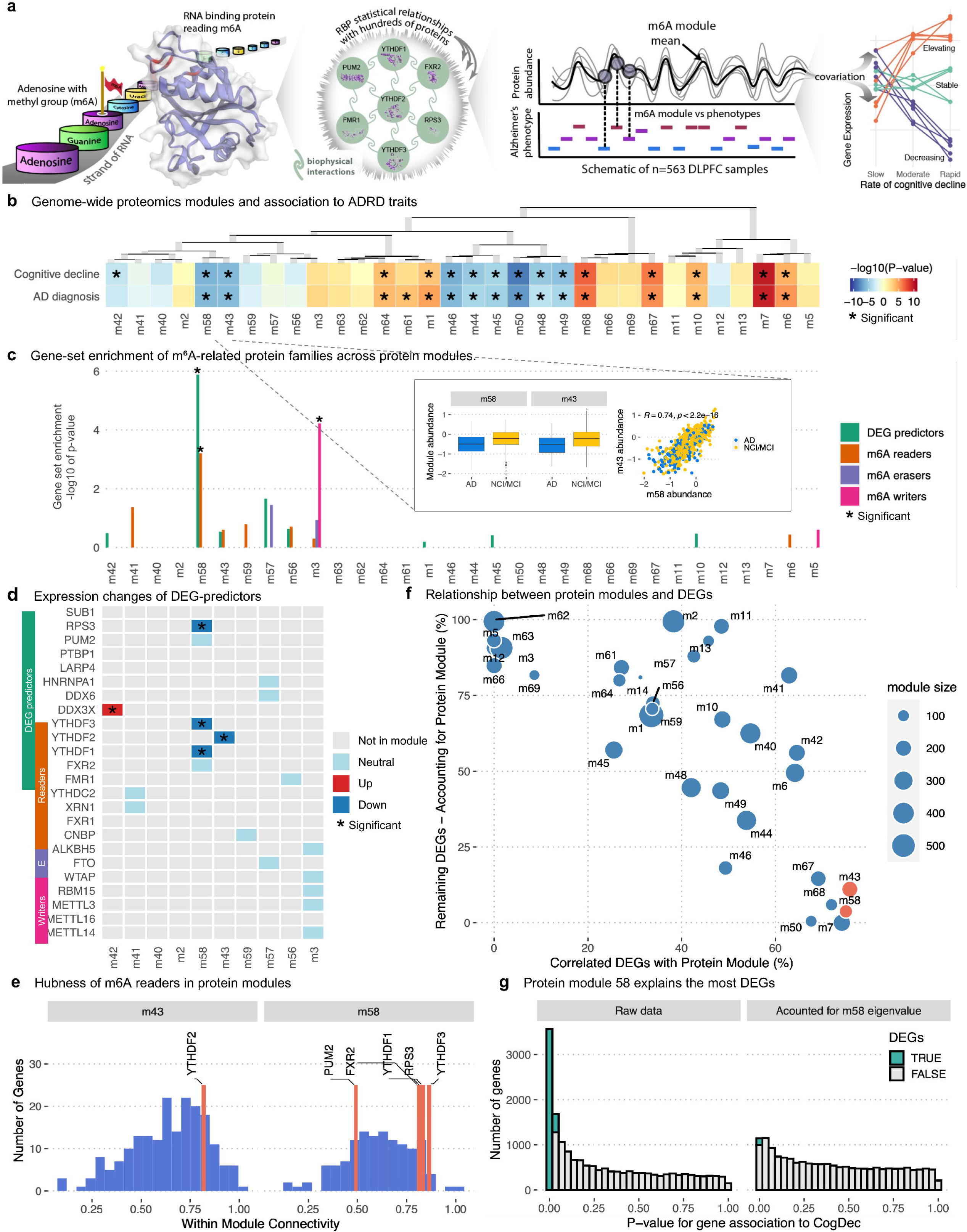
Protein modules of m6A readers explain most of the DEGs

At the individual protein level, the expression of RPS3, YTHDF1, and YTHDF3 in m58, and YTHDF2 in m43 (194 proteins), was reduced in AD individuals with rapid cognitive decline (**Fig. 2d**). Confirming these results we replicated in an independent TMT proteomics dataset (n=288), and meta-analysis across both datasets confirmed significant downregulation of these RBPs (RPS3: p=4.4×10^−9^, YTHDF1: p=1.5×10⁻□, YTHDF2: p=7.6×10⁻□, YTHDF3: p=2.9×10⁻□) and their containing modules (m58: p=1.0×10^−14^, m43: p=4.6×10^−12^) (**Supplementary Fig. 2a**). Notably, m58 and m43 were highly correlated with each other, and they were negatively associated with AD and cognitive decline (**Fig. 2b,c**). We did not detect those m6A reader changes at the RNA level and the preservation of the m58 and m43 modules at the RNA level (**Supplementary Fig. 2b,c**). This observation aligns with our previous study, which showed a generally poor correlation between RNA and protein levels and their covariance structures in human brains^12^. To evaluate whether these m6A readers connect with other proteins in the same module, we calculated the strength of intra-module connectivity of each protein within m58 and m43. Indeed, the m6A readers demonstrated high connectivity with other proteins in their respective modules, indicating their “hub” function (**Fig. 2e**).

This led us to hypothesize that if m6A readers control the AD gene signature, m58 and m43 expression would account for the DEGs associated with cognitive decline (**Fig. 2a**). To test this hypothesis, we leveraged the paired proteomic and transcriptome data from 549 DLPFC samples. Specifically, we computed two key metrics: the percentage of DEGs significantly correlated with protein module levels, and the percentage of DEGs remaining after accounting for module effects. Strikingly, both metrics revealed the substantial impact of m58 and m43 on the DEGs associated with cognitive decline (**Fig. 2f** and **Supplementary Table 6**). The initial set of 3,971 DEGs was dramatically reduced to 149 genes after accounting for the effects of m58 (**Fig. 2g**), and similar results were observed for m43 (**Supplementary Fig. 2d)**. These results strongly suggest a pivotal role of m6A reader modules in shaping the cognitive decline gene signature in AD.

Taken together, we show that m6A readers and their broader mechanisms are negatively associated with cognitive decline and AD diagnosis. These m6A readers interact broadly with other proteins within the same module. Removing the effect of m6A reader modules offsets most of the gene expression changes related to cognitive decline. Importantly, these results support the predictions of our model, indicating the critical roles of m6A-related RBPs in shaping brain gene signatures in AD dementia.

### Genetic perturbations of m6A-related RBPs recapitulate the AD-associated gene signatures in iPSC-derived cortical cells

Consistent transcriptomic and proteomic evidence indicates that m6A-related RBPs are involved in shaping AD gene signatures. However, whether they play a causal role or are merely correlates remains unclear. Because m6A readers were downregulated in AD brains (**Fig. 2d**), we reasoned that reducing their expression in human cells would help define their functional consequences. To test this, we used a commercially available neural progenitor cell (NPC) line derived from standard iPSC cortical differentiation, representing an early cortical progenitor stage as confirmed by established marker expression^13–15^.

To enable CRISPR/Cas9 editing, we transduced NPCs with Cas9-expressing lentiviruses and expanded the cultures for two weeks (**Supplementary Fig. 3a**). This strategy introduced CRISPR-Cas9 machinery while allowing the cells to mature toward a late progenitor stage. We then cultured the Cas9-expressing cells for an additional 6 days before transfecting them with synthetic guide RNAs (gRNAs) targeting major m6A readers (YTHDF1, YTHDF2, YTHDF3, and FMR1), writers (METTL14, METTL3, and WTAP), or the eraser FTO. After a further 9-day perturbation period, the cultures exhibited the expected heterogeneity of cortical differentiation, comprising PAX6^+^ progenitor cells and MAP2^+^ neurons (**Supplementary Fig. 3b)**. We harvested both RBP-targeting and non-targeting control (NTC) cultures for RNA-seq and knockdown-validation experiments. Editing efficiency was confirmed at the DNA level using a T7 endonuclease I assay, and knockdown efficiency was further verified using western blotting and RNA-seq–derived transcript abundance (**Fig. 3b** and **Supplementary Fig. 3c,d**).

**Figure 3.**
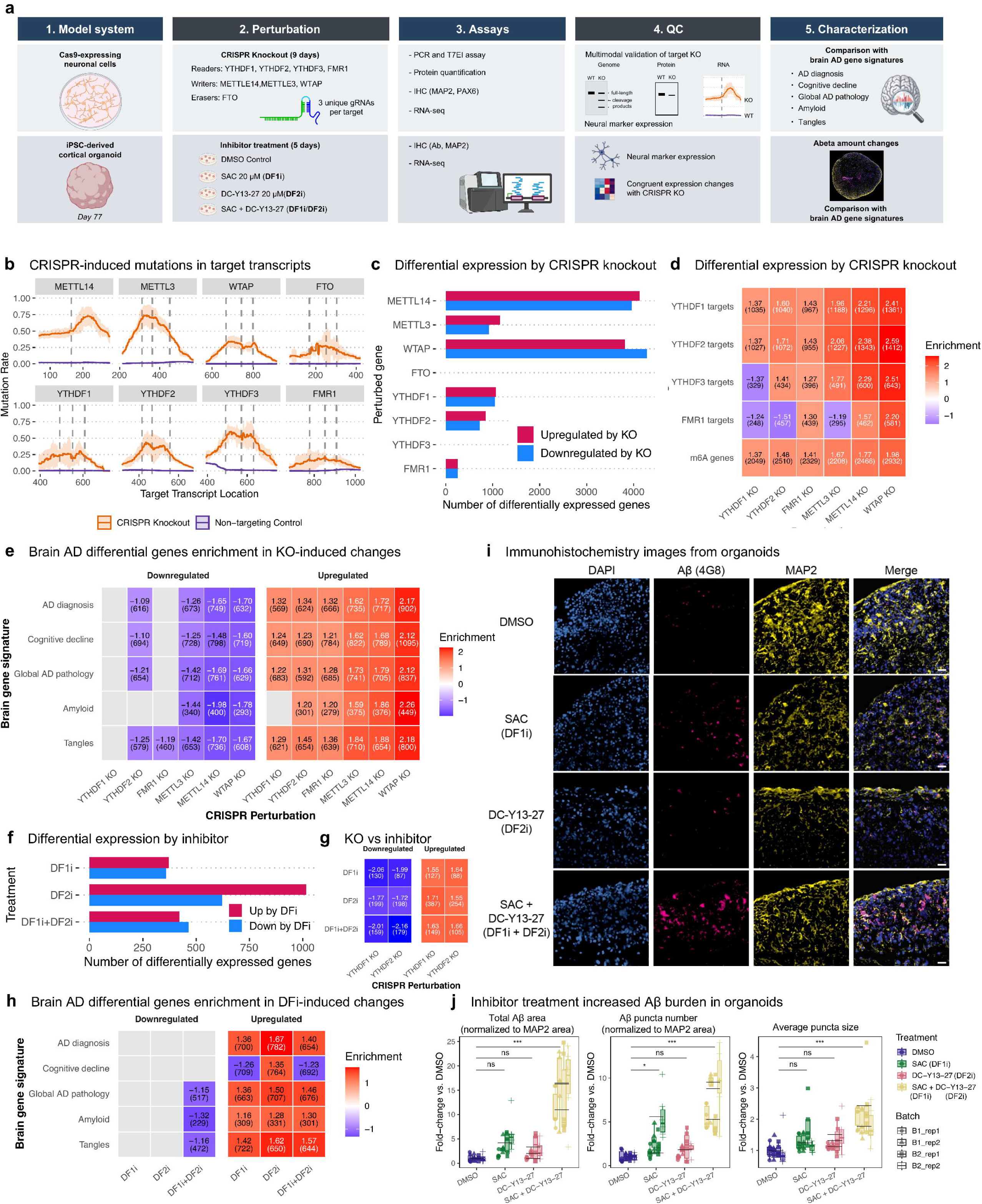
Perturbing m6A-associated RBPs in neuronal cells recapitulates the AD brain transcriptome states

We next conducted differential gene expression analysis on the RNA-seq data to characterize KO-induced transcriptional changes. This analysis revealed hundreds to thousands of DEGs for each perturbation, with the exception of YTHDF3 and FTO (**Fig. 3c** and **Supplementary Table 7**). To determine whether these KOs affect genes bound by m6A readers or genes harboring m6A modifications, we performed gene-set enrichment analysis. We found that these KOs significantly upregulated corresponding target genes and genes with known m6A modifications, consistent with these RBPs functioning as post-transcriptional suppressors (**Fig. 3d)**. We next examined whether the KO-induced DEGs overlapped with five AD-associated gene signatures derived from postmortem DLPFC tissue, specifically those linked to AD diagnosis, cognitive decline, global AD pathology, amyloid burden, and tau pathology. Across all five AD-associated gene signatures, we observed a strong and consistent positive correlation, whereby KO-induced upregulated genes correspond to AD-upregulated signatures, and KO-induced downregulated genes mapped to AD-downregulated signatures (**Fig. 3e** and **Supplementary Table 8)**. These results demonstrate that reducing YTHDFs activity is sufficient to recapitulate key AD transcriptional states in cultured iPSC-derived cortical cells.

### Pharmacological inhibitions of YTHDF proteins recapitulate AD-related gene signatures and A**β** pathology in iPSC-derived cortical organoids

To validate the CRISPR-based perturbation findings in an orthogonal 3D human neural model, we generated cortical organoids from iPSC lines derived from cognitively normal ROSMAP participants using established differentiation protocols^16^. Day-77 organoids were treated with SAC (YTHDF1 inhibitor)^17^, DC-Y13-27 (YTHDF2 inhibitor)^18^, or both inhibitors in combination, followed by multimodal characterization five days later, including RNA-seq and immunostaining (**Fig. 3a**). To confirm the expected 3D organization, we stained organoids for PAX6/TBR2 (progenitor zones) and TBR1/CTIP2/SATB2 (neuronal layers) (**Supplementary Fig. 3e**).

RNA-seq analysis revealed hundreds of DEGs across all inhibitor conditions (**Fig. 3f** and **Supplementary Table 9**). Notably, transcriptomic changes induced by inhibitor treatment were strongly concordant with those observed in the corresponding CRISPR KOs (**Fig. 3g**), indicating on-target engagement of YTHDF1/2. We next evaluated whether inhibitor-induced DEGs aligned with the same five AD-related transcriptomic signatures. Across all signatures, AD-upregulated genes showed strong and consistent enrichment among inhibitor-upregulated genes, whereas enrichment of AD-downregulated genes was more limited and primarily detectable in the DF1+DF2 inhibition condition (**Fig. 3h**). Thus, pharmacological inhibition preferentially recapitulates the AD-upregulated gene program.

Because Aβ predominantly accumulates within neuronal regions, Aβ measurements were normalized to MAP2+ area to account for regional and organoid-to-organoid variability. Organoids treated with either SAC or DC-Y13-27 alone showed minimal changes in total Aβ area, Aβ puncta number, or average puncta size, whereas combined inhibition of YTHDF1 and YTHDF2 led to marked increases across all Aβ metrics compared with DMSO controls (**Fig. 3i-j; Supplementary Fig. 3f-g**).

Together, genetic and pharmacological perturbations converge to demonstrate that reduced YTHDF activity is a causal driver of both AD-associated transcriptomic signatures and AD-like neuropathological phenotypes.

### Partial overlapping of m6A reader binding sites with m6A-modification sites in postmortem DLPFC

In order to examine whether m6A readers recognize m6A-modified transcripts in postmortem human brains, we conducted enhanced crosslinking and immunoprecipitation (eCLIP) to profile transcriptome-wide binding sites for YTHDF1, YTHDF2, YTHDF3, and FMR1, along with m6A-modified sites on DLPFC samples from 39 AD and 42 non-cognitively impaired (NCI) individuals (**Supplementary Table 10**). First, we performed QC analysis to assess the variability across sequencing batches and between samples to ensure the consistency and reliability of the data (**Supplementary Fig. 4a,b**). We then used the peak-calling tool CLIPper to identify peaks and determined consensus peaks, i.e., peaks present in at least 50% of samples, for YTHDF1, YTHDF2, YTHDF3, FMR1, and m6A (**Supplementary Fig. 4c** and **Supplementary Table 11**). Consistent with previous reports^19,20^, peaks for YTHDF1, YTHDF2, YTHDF3, and FMR1 were primarily located both at the boundary between coding sequences (CDS) and 3′ untranslated regions (3′ UTRs) as well as within the 3′ UTRs, whereas m6A peaks were primarily found at the CDS-3′ UTR boundary (**Fig. 4a**).

**Figure 4.**
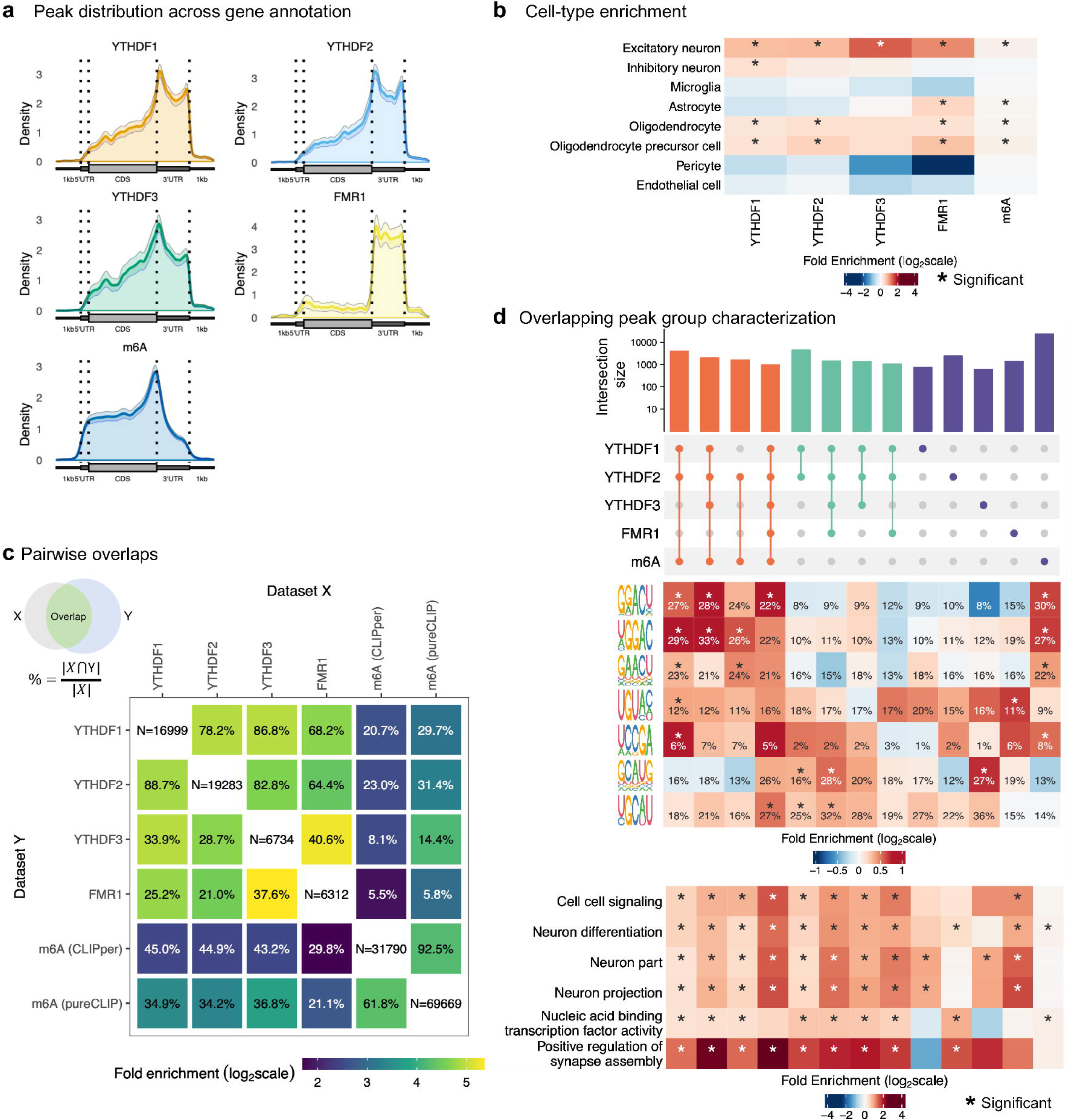
Characterization of m6A reader bindings and m6A modifications in DLPFC from older adults

To understand in which cell type(s) the m6A modifications and YTHDF binding activities occur, we tested whether genes containing m6A reader and m6A peaks were enriched among marker genes of major brain cell types. We found that the genes with m6A reader and m6A peaks were primarily enriched in excitatory neurons; genes with YTHDF1, YTHDF2, FMR1, and m6A peaks were also enriched in oligodendrocytes and oligodendrocyte precursor cells; genes with m6A and FMR1 peaks were also enriched in astrocytes (**Fig. 4b**).

It is well known that m6A readers recognize m6A sites and mediate downstream RNA metabolism^21^. We reasoned that for this canonical pathway, we would expect to observe overlapping peaks between m6A and m6A readers. Conversely, a lack of overlapping peaks between m6A and m6A readers would suggest a non-canonical m6A-independent pathway. To this end, we first examined the peak overlaps for any given pair among m6A readers and m6A. As expected, we detected a significant overlap between m6A readers and m6A (3.26- to 17.9-fold). In addition, we also observed a much stronger overlap among m6A readers (21.1- to 40.8-fold) (**Fig. 4c**). However, the maximum observed overlap between m6A reader peaks and m6A peaks was 45%, suggesting that m6A readers can bind to RNAs independently of m6A sites.

To further examine sequence dependency, we conducted motif analysis on YTHDFs and m6A peaks. As expected, we found that the GGACU motif, a common DRACH sequence where the m6A modifications occur on RNA, was enriched in the peaks for YTHDF1, YTHDF2, YTHDF3, and m6A (**Supplementary Fig. 5a**). Notably, we also detected non-canonical motifs, where different nucleotides flanked the A/m6A for reader and m6A peaks (**Supplementary Fig. 5a**). We then clustered all peaks into three major groups based on the peak colocalization pattern: Peak Group 1 with colocalized peaks for m6A and YTHDF1, YTHDF2, YTHDF3, or FMR1; Peak Group 2 with colocalized peaks for RBPs only; and Peak Group 3 with no colocalized peaks (singletons) (**Fig. 4d** and **Supplementary Table 11**). Interestingly, both canonical and non-canonical m6A/A motifs were significantly enriched in Peak Group 1, but to a lesser extent in Peak Groups 2 and 3 (**Fig. 4d** and **Supplementary Fig. 5b**). Despite these differences, the genes bound by canonical or non-canonical pathways largely overlap. Specifically, of the 3,927 genes with peaks in Group 1 and the 2,846 genes in Group 2, 2,325 genes (81.7%) overlapped. This considerable overlap suggests that the co-occurrence of both peak types within the same gene is a prevalent feature. Reflecting this observation, both groups were strongly associated with synaptic signaling assembly, indicating their convergent functional effects (**Fig. 4d**).

For additional confirmation from a different methodology, we also used pureCLIP to identify m6A modification sites at a single-nucleotide resolution and detected similar peak distribution and motif enrichment to those revealed by CLIPper (**Supplementary Fig. 5c,d**). Taken together, our results suggest that the m6A readers may regulate gene expression post-transcriptionally via both m6A-dependent canonical and m6A-independent non-canonical pathways^22^. Our findings suggest a broader scope for the gene regulatory potential of m6A reader proteins.

### Altered m6A modifications and m6A-reader binding activities in AD brains

We investigated differential peaks for m6A modification and YTHDF binding activities in AD versus NCI brains. First, we took a binary approach to call the presence or absence of peaks for any given location in the transcriptome, and then calculated the overall peak numbers in NCI and AD brains. After adjusting for sequencing depth, batch, and RNA integrity, we detected a general reduction of peaks for all reader proteins and m6A in AD brains (**Fig. 5a**). Interestingly, the reduction in peaks for m6A reader proteins occurred primarily in 3’ UTRs, whereas the reduction in peaks for m6A was in CDS (**Fig. 5a**). Along with the reduced expression of YTHDFs and YTHDF-containing protein modules in AD brains (**Fig. 2b,c,d**), our results support a reduced trend of m6A modification and m6A reader binding activities in AD brains, which is largely in line with the results from previous animal and human studies^23–25^.

**Figure 5.**
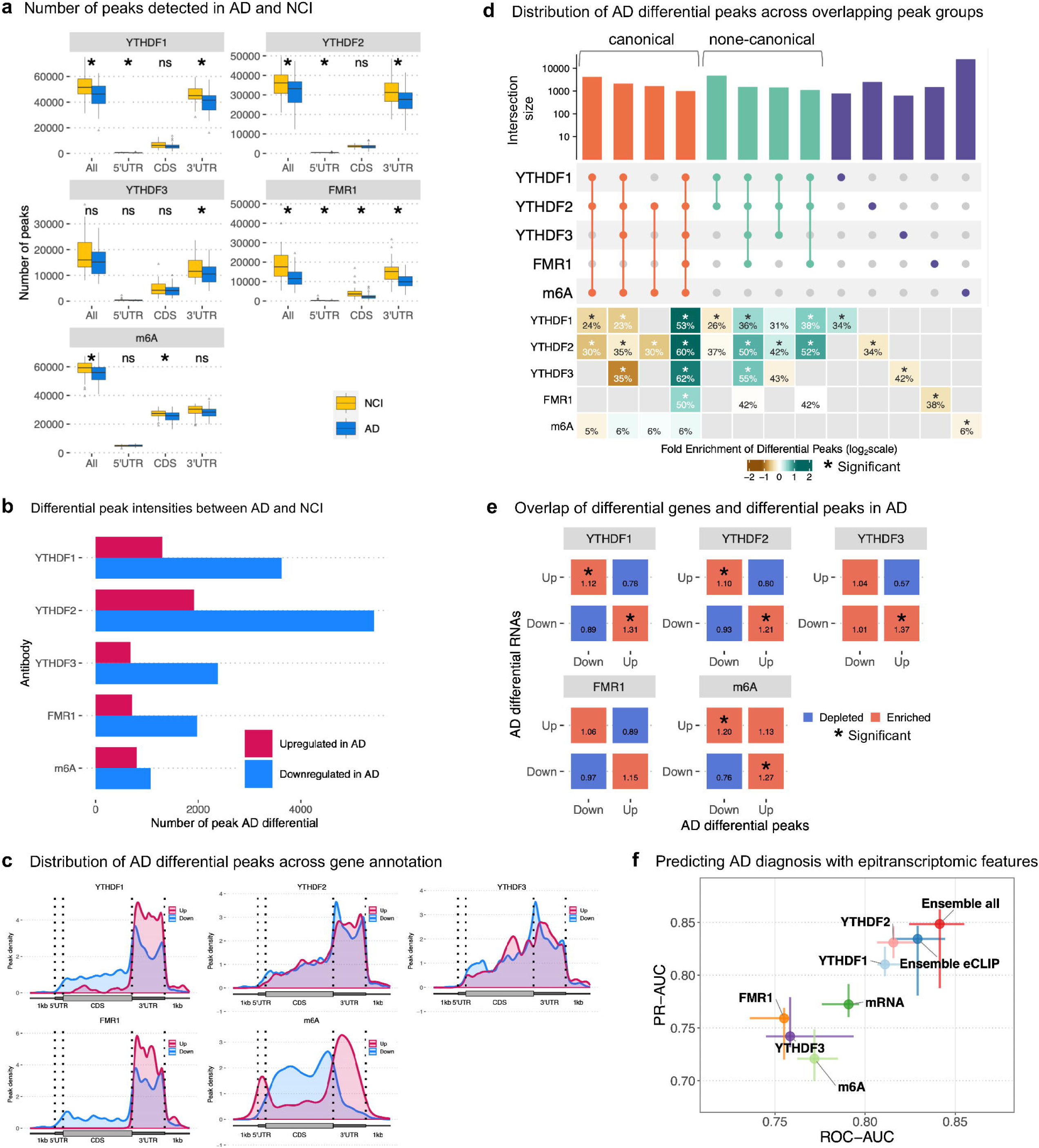
Differential m6A-reader bindings and m6A modifications in AD brains

For robustness, we also used a graded approach to gain a deeper understanding of differential m6A reader binding activities and m6A modifications in AD. To achieve this, we counted read numbers spanning each consensus peak and then normalized the reads to the parental RNA expression to derive a normalized peak intensity. This graded approach enabled transcriptome-wide peak association analysis, identifying peaks with significant differential intensity in AD. We detected more up- and down-regulated differential peak intensities in m6A reader proteins than in m6A modifications (**Fig. 5b** and **Supplementary Table 12**). Interestingly, the up-and down-regulated peaks for m6A reader proteins were primarily located in the 3’UTR, consistent with their typical peak distribution pattern (**Fig. 4a** and **Fig. 5c**). Conversely, the up-regulated m6A peaks were found in the 5’ UTR and 3’ UTR, whereas down-regulated m6A peaks were in the CDS, which deviated from the normal peak distribution pattern of m6A (**Fig. 4a** and **Fig. 5c**). This pattern shift of m6A differential peak intensity may reflect broader RNA processing changes in response to AD pathogenic processes. Building on our previous clustering of peaks into three major groups based on their colocalization patterns (**Fig. 4d**), where Peak Group 1 represents the m6A-dependent canonical pathway and Peak Groups 2 and 3 represent the m6A-independent non-canonical pathway, we sought to examine the enrichment of AD differential peaks within these groups. Differential peak enrichment was observed in both m6A-dependent and independent pathways. Specifically, AD differential peaks were positively enriched in regions colocalized with all RBPs—YTHDF1, YTHDF2, YTHDF3, and FMR1—and m6A within Peak Groups 1 and most of Peak Groups 2 (**Fig. 5d**). Conversely, AD differential peaks were depleted in m6A-dependent pathways lacking FMR1 and in singleton peaks, with the exception of YTHDF1. These findings suggest that the m6A-independent non-canonical pathway may also play a prominent role in regulating AD gene signatures.

To examine whether the changes in m6A modification and m6A reader binding activity in AD brains are associated with transcriptomic changes, we mapped peak changes to individual DEGs. We detected an inverse correlation for all m6A readers, i.e., upregulated DEGs were associated with decreased peak intensities, whereas downregulated DEGs were associated with increased peak intensities (**Fig. 5e**). This inverse relationship is exemplified by genes showing significant differential expression and opposite-direction changes in m6A reader binding across multiple reader proteins (**Supplementary Fig. 5e**). Given the significant overlap of peaks between reader proteins (**Fig. 4c**), this consistent relationship across all reader proteins indicates that these proteins may work synergistically to regulate mRNA metabolism. The anti-correlation relationship between individual DEGs and peak intensities also suggests that reader protein binding activities may reduce RNA stability and/or increase RNA decay in AD brains, leading to reduced gene expression.

Finally, we developed predictive models for AD diagnosis using eCLIP and/or RNA-seq data (**Fig. 5f**). Notably, YTHDF2 demonstrated superior accuracy in predicting AD compared to other m6A readers, with its predictive power close to an ensemble model combining eCLIP and transcriptomic data (**Fig. 5f**). Along with our proteomics and gene perturbation results, our findings indicate that YTHDFs, especially YTHDF2, are significantly associated with AD pathology and provide valuable insights into the molecular mechanisms underlying gene expression changes in AD brains.

## Discussion

In this study, we uncover a novel mechanism underlying molecular changes in AD, highlighting the pivotal roles of m6A reader proteins in regulating AD gene signatures. Utilizing an explainable machine learning framework, we identify m6A-related RBPs as key regulators of gene expression changes in AD. Importantly, we then validate our predictions across multiple independent approaches, including proteomic analysis, eCLIP-seq experiments, and *in vitro* perturbations. Specifically, we demonstrate that YTHDFs and YTHDF-containing protein modules are downregulated in AD; ablation of YTHDF-containing modules *in silico* accounts for the majority of differentially expressed genes in AD; reducing m6A-related RBP activity in iPSC-derived neuronal cells and 3D organoids recapitulates the AD-associated gene signatures and Aβ-associated neuropathological phenotypes; m6A reader binding activities and m6A modifications are altered in AD brains. Our findings, supported by *in vivo* and *in vitro* evidence, position m6A readers as central regulators of the AD transcriptome and highlight their critical roles in AD pathogenesis.

While our model achieved an overall correlation of R=0.38 with differentially expressed genes (**Fig. 1c**), this modest correlation reflects inherent limitations in predicting complex transcriptional changes in Alzheimer’s disease. Subsampling analyses between independent RNA-seq batches revealed that correlation plateaus at approximately R≈0.63 even with large sample sizes, establishing an empirical upper bound constrained by biological variability and technical noise (**Supplementary Fig. 1b**). Our model’s performance thus captures approximately 60% of the empirically attainable signal. Importantly, despite this modest overall correlation, the ranking of key regulatory features remains highly consistent across independent datasets (Spearman r=0.77), with m6A readers consistently emerging as top predictors (**Supplementary Fig. 1d**). This consistency, coupled with our orthogonal experimental validations through proteomics, CRISPR perturbations, and eCLIP experiments, provides strong evidence for the biological significance of our findings. Additionally, prediction accuracy is fundamentally constrained by the limited availability of comprehensive RNA-protein interaction data^8^, as global RNA-binding profiles exist for only approximately 10% of known RBPs, affecting all gene regulation-based models in this domain.

Numerous studies have demonstrated that m6A pathways play essential roles in neural development, synaptic plasticity, stress response, learning, and memory^26–28^. Emerging evidence also reveals that disruption of m6A pathways contributes to neurodegeneration and cognitive decline. For instance, the levels of m6A writers, m6A readers, and global m6A modifications are altered in AD mouse models and human AD brains^23–25,29–31^. Furthermore, reduced neuronal m6A modifications lead to significant memory deficits, extensive synaptic loss, and neuronal death in mouse models^23,24^. The m6A readers, such as YTHDF1 and YTHDF2, have been shown to play essential roles in learning and memory in animal models by regulating local protein synthesis and controlling mRNA localization^20,32,33^. Our study expands upon previous research by providing solid evidence for the pivotal role of m6A reader proteins in regulating AD gene signatures. Using AI-based integrative analysis of human brain multi-omics data, along with gene perturbation experiments in human neuronal cells, we discover the causal roles of YTHDFs in shaping AD gene signatures. This integrative and unbiased approach presents a robust framework for identifying, *in silico* testing, and experimentally validating gene regulators. To further explore the potential causative contribution of YTHDF proteins to AD, we analyzed the most recent multi-ancestry Alzheimer’s disease GWAS meta-analysis (111,326 cases and 677,663 controls)^34^. We observed a nominally significant gene-level association for YTHDF1 (Cauchy p = 0.0097), while YTHDF2 and YTHDF3 showed no association (p ≈ 0.80 and 0.70, respectively). FMR1, located on the X chromosome, lacks available GWAS summary statistics. However, FXR2, an autosomal homolog of FMR1 that shares functional domains and is also highlighted by our DEcode-tree model (**Fig. 1g**) and involved in the protein module 58, shows significant association (Cauchy p = 0.0019). Loss-of-function constraint metrics from gnomAD^35^ reveal that these genes are highly dosage-sensitive, with LOEUF scores (where <0.35 indicates strong constraint) of: 0.515 for YTHDF1 (moderate constraint), 0.132 for YTHDF2 (very strong constraint), 0.272 for YTHDF3 (strong constraint), and 0.321 for FXR2 (strong constraint). The extreme constraint on YTHDF2 and YTHDF3 may explain their lack of strong GWAS signals, as highly constrained genes often contribute to disease risk through subtle regulatory variants rather than common coding changes.

These results provide a holistic view of the m6A epi-transcriptomic landscape of the aging brain, through comprehensive epi-transcriptomic profiling of m6A modifications and m6A reader binding in human brains. The functions of YTHDFs in regulating RNA metabolism have been controversial. The prevailing model suggests that each YTHDF protein has distinct functions by binding to specific mRNAs, e.g., YTHDF1 enhances mRNA translation, while YTHDF2 mediates mRNA decay^21^. Conversely, the competing model proposes that YTHDF proteins have redundant functions, binding identically to all m6A sites on mRNAs and together mediating mRNA degradation in HeLa cells^36^. Furthermore, recent evidence supports a context-dependent model where YTHDFs interact with various partner proteins and RNAs within cytosolic condensates, such as stress granules^22,37–39^. Our findings align with both the competing model and the context-dependent model for the following reasons. First, we observed significant overlap among m6A reader binding sites, either m6A-dependent or m6A-independent, indicating that different m6A reader proteins can bind the same sites on the same transcript; alternatively, different transcripts of the same gene can have the same sites recognized by different readers. Second, we showed a consistent inverse correlation between each reader and AD DEGs, implying the synergistic effect of reader proteins on mRNA degradation. Moreover, our results support the context-dependent model. For instance, we detected both canonical, m6A-dependent motifs (Peak Group 1) and non-canonical m6A-independent motifs (Peak Group 2) (**Fig. 5d**). This suggests that reader proteins may bind directly to non-canonical sequences, regardless of m6A modification, or indirectly via protein-protein interactions. Genes bound via both canonical and non-canonical pathways are strongly associated with synaptic signaling and neuronal projection (**Fig. 4d**), indicating their convergent cellular functions. In summary, our results reveal the complexity of YTHDF proteins in gene regulation, which may allow for fine-tuned, context-dependent gene regulations in AD development.

Although we employed an integrative framework for gene regulator nomination and validation along with extensive multi-omics datasets, a few limitations should be noted. First, the ROSMAP cohort mainly consists of white individuals, limiting the generalizability of our findings to other racial and ancestral groups. Second, while this constitutes a large eCLIP dataset, this sample size is not large enough to extend beyond a case-control design, which would likely require several hundred samples, based on other omics data acquired in this same cohort. Our findings highlight m6A readers, especially YTHDF2, as potentially important indicators of AD pathology. However, we acknowledge important limitations regarding disease specificity. Comparing these m6A-mediated mechanisms across different neurological conditions would be valuable future work to determine the specificity of YTHDF dysregulation in AD versus other neurodegenerative diseases. Third, key m6A-associated RBPs, although supported by experimental validation, may represent only a fraction of the RBPs that regulate AD gene expression. Furthermore, we focus primarily on m6A modifications, but other RNA modifications, such as N1-methyladenosine (m1A), could also be recognized by YTHDFs^40^. Fourth, our bulk RNA-seq approach may underrepresent regulatory changes in less abundant cell types like microglia, which play critical roles in AD pathophysiology. Single-nucleus RNA-seq data would provide clearer insights into cell-type-specific regulatory mechanisms and would be valuable for future applications of our framework. Thus, a worst-case interpretation of our results is that we have found a minimal set of proteins that account for human brain changes, which can reproduce genome-wide signatures of disease in human model systems. This is as powerful a conclusion as can be drawn from primary data and early-stage validation tests. In terms of its implications even beyond Alzheimer’s disease, this study provides a robust and reliable interpretation of AD gene signatures in human brains, opening new avenues for harnessing integrative research strategies to uncover the top regulators of global expression.

## Methods

### Participants

Participants came from ROS or MAP studies. The two studies began enrollment in 1994 and 1997, respectively^41^. ROS and MAP are conducted by the same team of investigators and share a large common core of measures, which allows combining the data for joint analyses. Participants enrolled without known dementia and agree to annual detailed clinical and cognitive evaluation, as well as brain donation after death. Both studies were approved by an institutional review board of Rush University Medical Center. At enrollment, each participant signed informed consent and an Anatomical Gift Act.

### Cognitive and clinical evaluations

AD diagnosis was based on criteria of the joint working group of the National Institute of Neurological and Communicative Disorders and Stroke and the Alzheimer’s Disease and Related Disorders Association (NINCDS/ADRDA) as previously reported^42^. Uniform structured cognitive and clinical evaluations are administered each year by examiners blinded to data from prior years. The cognitive battery contains a total of 21 cognitive performance tests. Briefly, 11 tests informed on diagnosis of AD dementia, mild cognitive impairment, and no cognitive impairment^42–44^. Further, 19 tests are used to construct a global composite measure of cognitive function^45^. The longitudinal rate of decline was computed for each participant using linear mixed models with adjustment for the effects of age, sex, and education, which estimate person-specific slopes. Additional models also controlled for common brain pathologies, resulting in person-specific residual decline as an estimate of resilience^46^.

### Neuropathological examination

Tissue was dissected from 8 brain regions to quantify the burden of parenchymal deposition of beta-amyloid and the density of abnormally phosphorylated paired helical filament tau (PHFtau)-positive neurofibrillary tangles. Tissue sections (20µm) were stained with antibodies against beta-amyloid protein and PHFtau protein, and quantified using image analysis and stereology, as previously described^46,47^. In addition, the assessment captured cortical Lewy bodies, hippocampal sclerosis, TDP-43, macro- and micro-infarcts, as well as atherosclerosis and arteriolosclerosis^46,47^.

### RNA-seq gene expression

ROSMAP RNA-seq data of DLPFC regions were downloaded from the AMP-AD platform (syn3388564) and processed as previously described^12^. Briefly, paired-end RNA-Seq data were first aligned by STAR v2.6^48^ to the human reference genome (Release 27 GRCh38). Picard tools were used to assess the quality of the aligned bam files. Transcript raw counts were calculated by Kallisto (v0.46)^49^. Samples were excluded if the total mapped reads were less than 5 million. A total of 17,294 mRNAs were expressed in >50% of the sample with at least 10 counts in each sample. These genes were included in the downstream normalization process. First, the CQN (conditional quantile normalization) was applied to adjust a sequence bias from GC content and gene length^50^. Next, the adjusted gene counts matrix was converted to log2-CPM (counts per million), followed by quantile normalization using the voom function implemented in the limma R package^51^. Finally, a linear regression model was applied to remove major technical confounding factors, including post-mortem interval, sequencing batch, RQN (RNA quality number), total spliced reads reported by STAR aligner, and metrics reported by Picard and Kallisto.

### Mass spectrometry-based proteomics using isobaric TMT

We downloaded ROSMAP TMT proteomics data from frozen tissue of the DLPFC from the Synapse repository (syn30390636 and syn2580853). The detailed experimental procedure can be found in the description of the repository and the previous reports^52,53^. Briefly, digested protein samples were labeled with isobaric TMT and fractionated using high-pH liquid chromatography. Fractions were then analyzed by LC-MS (liquid chromatography-mass spectrometry). The resulting MS spectra were searched against the Uniprot human protein database and quantified. The effects of the experimental batch, PMI, sex, and age at death on quantified protein abundance were regressed out using linear regression. Proteins with missing values in more than 50% of subjects were excluded. A total of 7,649 proteins in 596 persons passed the final QC. For replication analysis of protein-cognitive decline relationships, we utilized an independent TMT proteomics dataset (n=288) from DLPFC regions generated through the AMP-AD project (syn51732482), encompassing multiple cohorts (ROSMAP, Minority Aging Research Study^54^, and African American Clinical Core^55^) conducted by the Rush Alzheimer’s Disease Center with harmonized cognitive phenotyping and identical TMT protocols. In addition to the standard covariates, racial background was included as a covariate in the replication analysis.

### Transcriptome/proteome-wide association study

Associations between transcript or protein levels and continuous or categorical outcomes were tested using linear regression models implemented in the limma package (v3.50.3) in R^51^. Age at death, sex, and years of education were used as covariates to control for potential confounding factors.

### DEcode-tree model

We calculated the median transcript length and median intron length for each gene across all annotated transcripts to represent gene structures while accounting for transcript variability. To represent exon counts per gene, we computed a weighted average of exon numbers, with each transcript’s exon count weighted by its length, giving greater influence to longer transcripts. Furthermore, we calculated the weighted average GC content for the 3’ UTR, 5’ UTR, exon, and intron regions of each gene. For each region, the GC content of each transcript was weighted by its proportion of the total length of that region across all transcripts of the gene. For RNA and DNA features, we downloaded the genomic locations of binding sites for 172 RNA-binding proteins (RBPs) from POSTAR2 (as of October 2018), 217 microRNAs (miRNAs) from TargetScan Release 7.2, and 833 transcription factors (TFs) from the Gene Transcription Regulation Database (GTRD) (as of October 2018). Next, we mapped the binding sites of RBPs, miRNAs, and TFs to promoters and exons. Promoter regions for each gene were defined as the region spanning from 2,000 base pairs upstream to 1,000 base pairs downstream of the transcription start site (TSS). For cell-type-specific expression signatures, we utilized single-cell RNA sequencing (scRNA-seq) data from adult human brains^56^. Gene counts were quantified using Kallisto (v0.46) with annotations from GENCODE Release 27. Reads from the same cell type were aggregated to create pseudo-bulk datasets. We computed log2 counts per million (CPM) and calculated fold changes between cell types relative to the average expression across all cell types.

We trained predictive models using genomic and RNA features to estimate t-statistics for differential gene expression associated with cognitive decline and AD diagnosis. Extreme Gradient Boosting (XGBoost) regression models were implemented using the xgboost package (v1.7.7.1) within the R programming environment, interfaced with the mlr package (v2.19.1). The primary objective was to minimize the root mean square error (RMSE), with model performance evaluated based on RMSE and Pearson’s correlation coefficient between predicted and observed t-statistics. The model was configured with fixed parameters for the objective function, evaluation metric, number of boosting rounds, learning rate, booster type, number of processing threads, and regularization terms. Hyperparameter tuning was performed to optimize model performance. Adjusted hyperparameters included tree complexity parameters (maximum tree depth and minimum child weight), regularization strengths (L1 and L2 penalties), and subsampling ratios for both training instances and features. Hyperparameter optimization was conducted using systematic exploration within the mlr framework to identify the optimal model configuration based on the lowest RMSE and the highest correlation. To assess the model’s ability to predict differentially expressed genes (DEGs), we constructed receiver operating characteristic (ROC) curves by comparing predicted t-statistics to actual DEG status. The area under the ROC curve (AUC) was calculated to quantify classification performance. We estimated the AUC along with a 90% confidence interval from 100 bootstrap samples to account for variability in the model’s performance, providing a robust measure of predictive accuracy.

We employed SHapley Additive exPlanations (SHAP) to identify the most influential predictive features. Predictive features were normalized by capping values at the 99th percentile and scaling to [0,1], regulatory binding site counts were normalized by maximum values with features having fewer than 30 positive observations excluded, and cell-type-specific expression values were scaled without centering. SHAP contribution values were computed for each feature using XGBoost’s built-in TreeSHAP implementation across five trained models, with SHAP values calculated on respective test sets using exact computation. Feature importance was ranked by averaging absolute SHAP values across all models and test samples. Enrichment analysis of m6A readers and interacting proteins of YTHDF1, YTHDF2, and YTHDF3 was conducted using the fgsea package (v1.20) with default parameter settings. RBPs were ranked by absolute SHAP importance scores, with gene sets defined from known m6A readers and protein-protein interactions. Statistical significance was assessed using false discovery rate (FDR < 0.05).

We analyzed relationships between predictive features using SHAP value correlations. Pairwise Pearson correlations were calculated between SHAP values across all features for each model separately, and the final correlation matrix was constructed by averaging correlations across models, retaining only features with complete correlation profiles. The correlation matrix was converted to a distance matrix using the transformation d = 1 - r, where r represents the correlation coefficient, and Uniform Manifold Approximation and Projection (UMAP) was applied to this distance matrix using the umap R package with parameters: n_neighbors = 15, min_dist = 0.5, and random_state = 42 for reproducibility.

### Protein co-abundance module analysis

To identify protein modules statistically, we employed the consensus clustering approach implemented in SpeakEasy (v1.0.0)^57^. We computed the composite metric for each module by averaging the normalized levels of all proteins within the module for each individual. To assess whether the protein modules are preserved in the transcriptome, we used the modulePreservation function from the WGCNA R package (v1.72)^58^. The network type was set to ‘signed,’ and all other parameters were kept at their default settings.

### Protein module impact assessment on DEGs

To assess the influence of m6A reader protein modules on cognitive decline-related DEGs, we computed two key metrics: the percentage of DEGs correlated with protein module levels and the percentage of remaining DEGs after accounting for module effects. First, DEGs associated with cognitive decline were identified from RNA-seq data from 549 samples with paired TMT proteomic measurements using the limma package, adjusting for covariates such as age at death, sex, and years of education. This analysis identified a total of 3,971 DEGs at an FDR of 5%. To examine the relationship between the DEGs and protein modules, we performed Pearson correlation analysis between the expression of each DEG and the eigenvalue (module summary) of each protein module. We then calculated the percentage of DEGs that showed significant correlations with module levels at an FDR of 5%. Following this, we conducted a regression analysis for each protein module to determine the DEG count after adjusting for the module’s effects. Module eigenvalues were included as covariates in the limma model to assess the contribution of each protein module individually. Finally, we calculated the reduction in the number of significant DEGs after adjusting for the effects of these modules, with significance assessed at an FDR of 5%.

### Cell line generation and maintenance

We used a commercially available Neural progenitor cell (NPC) line derived from XCL-1 iPSCs from Stem Cell Technologies (200-0620). This NPC line was generated from standard cortical differentiation, equivalent to early NPCs, based on previously published protocols from us and others^13–15,59^. The NPCs were further matured into late NPCs by culturing and expanding them in STEMdiff Neural Progenitor Medium (StemCell Technologies, 05833) for 2 weeks. To establish stable Cas9 expression, early NPCs were transduced with lentiviruses carrying CRISPR Sp Cas9 pRCCH-CMV-Cas9-2A-Hygro virus at an MOI of 0.3 (Cellecta, SVC9-VS). After 48 hr, cells were selected with 100 μg/ml Hygromycin B (HygroB; Gibco, 10687010), cultured for 2 weeks to mature into late NPCs, and then banked and cryopreserved.

### Synthetic gRNA transfection

Synthetic guide RNAs targeting 8 m6A-relevant genes or 2 non-targeting (NT) controls were designed by and ordered from Synthego. Guide RNAs were provided as a lyophilized pool of two or three gRNAs, each targeting distinct sequences in exon 1 of the target gene. Each synthetic guide pool was resuspended in water to a 50 μM stock concentration. To perturb m6A-related RBPs, we first thawed and cultured the Cas9-expressing late NPCs for 6 days, then proceeded with transfections as described below. Cas9-expressing late NPCs were detached from dishes using Accutase, centrifuged at 300 x g for 4 min, then resuspended in pre-warmed Neural Progenitor Medium (without HygroB) containing 20 μM Rho-kinase inhibitor (Y-27632; StemCell Tech 72304). Cells were counted and diluted to 1 × 10^6^ viable cells/ml. Meanwhile, gRNAs were diluted to 200 nM in sterile 1X TE buffer, pH 8.0 (220 μl each). In a 15-ml conical tube, 10 ml pre-warmed Opti-MEM media (Gibco; Thermo 31985070) was combined with 60 μl Lipofectamine RNAiMax (Thermo Fisher 13778150) and mixed well by inverting. For each KO target, 220 μl of diluted gRNA was combined with 800 μl Opti-MEM/RNAiMax mixture, pipette-mixed, and incubated at RT for at least 5 min but not more than 30 min. For transfection, the prepared cell suspension was plated onto pre-coated Matrigel plates (1 × 10^6^ cells in 1 ml for 6-well; 5 × 10^4^ cells in 50 μl for 96-well) and gRNA/Opti-MEM/RNAiMax mixture was immediately added to each well (950 μl for 6-well; 50 μl for 96-well), with a final concentration of 10 μM Y-27632 and 20 nM gRNA. Additional wells were subjected to mock transfection (using sterile TE buffer in place of gRNAs). Six hours after transfection, the media was removed and replaced with fresh Neural Progenitor Medium (without HygroB). Full media changes were performed again at 2-, 3-, and 4-days post-transfection (dpt) with fresh Neural Progenitor Medium (+ 100 μg/ml HygroB). At 5 dpt, genomic DNA (gDNA) was isolated from 96-well plates (see below), and cells in 6-well plates were passaged (using Accutase) to additional wells of matrigel-coated 6-well plates for further expansion, with daily media changes until 9 dpt. This protocol was repeated 2 times for a total of 3 biological replicates. The resulting cultures were then used for RNAseq and immunofluorescence staining. For staining, cells were fixed in 4% PFA for 15 min, blocked and permeabilized in 10% normal goat serum with 0.1% Triton X-100. The cells were stained with primary antibodies against MAP2 (Novus Biologicals NBP1-40606) and PAX6 (DSHB AB_528427), followed by species-specific Alexa Fluor-conjugated secondary antibodies (Thermo Fisher Scientific).

### Genomic DNA isolation and PCR

At 5 days post-transfection (dpt), media was aspirated and discarded from each well, then 100 μl of ice cold QuickExtract reagent (LGC Biosearch Technologies QE0905T) was added to each well. The plate was incubated at 65°C for 6 min with 700 rpm shaking on a thermoshaker. Then 100 μl of lysate per well was transferred to a 96-well PCR plate. The plate was sealed, then incubated at 98°C for 2 min followed by a 4°C hold. Due to the high viscosity of isolated DNA, samples were diluted 4-fold in fresh QuickExtract reagent to enable accurate pipetting of low volumes. Undiluted and diluted gDNA were stored at -80°C. PCR reactions were set up on ice using AmpliTaq Gold 360 polymerase (Applied Biosystems; Thermo Fisher 4398881) with the following components: 25 µl of 2X Master Mix, 0.5 µl of 10 mM forward primer, 0.5 µl of 10 mM reverse primer, 5 µl of GC enhancer, 15 µl of water, and 4 µl of gDNA, to a total volume of 50 µl. Primers were designed to amplify a 600-1200 bp fragment surrounding the Cas9 cleavage site, ensuring at least 200 bp between the primer binding site and any gRNA target region. Lysates from mock-transfected/WT or KO cells were used as templates. PCR cycling conditions were set as follows: enzyme activation at 95°C for 10 seconds, followed by 40 cycles of denaturation at 95°C for 25 seconds, annealing at 52°C for 30 seconds (with the annealing temperature decreasing by 2°C for each of the first four cycles, starting at 60°C), and extension at 72°C for 1 minute. A final extension at 72°C for 6 minutes was performed, with samples held at 4°C.

### T7 Endonuclease I (T7EI) assay

T7EI assay was performed according to the manufacturer’s instructions (NEB M0302L). Briefly, a 2.5% agarose gel containing SYBRSafe dye (Invitrogen S33102; diluted 1:10,000) was prepared. 12.5 μl of each PCR product was mixed with 1.4 μl NEB Buffer 2. The PCR products were then denatured at 95°C for 5 min and re-annealed slowly (95-85°C: -2°C/sec; 85-25°C: -0.1°C/sec; 4°C hold). 0.5 μl of T7 Endonuclease I (T7EI) was added to each sample, followed by 15 min incubation at 37°C. 3 μl of 6X loading dye was then added to each T7-digested sample (WT and KO for each target), which were loaded to the agarose gel along with 100 bp ladder (Thermo Fisher SM0323) and run at 120V for approximately 1 hour. UV images were captured using the Azure 300 Chemiluminescent Imaging System (Azure Biosystems). Band intensities were quantified using Fiji. Editing efficiency was estimated using the standard formula: 100 x (1 – (1- fraction cleaved)^1/2^). The fraction cleaved was calculated by summing the intensities of the cleavage bands and dividing by the summed intensities of all bands (including cleaved and full-length). Intensities were normalized to the WT (unedited) sample so that the background intensity of cleavage bands = 0 for the WT.

### RNA isolation, library preparation, and sequencing

At 9 dpt, media was aspirated from two wells of a 6-well plate for each condition (8 KOs, 2 NT gRNAs). Cells were washed once with 1X DPBS, then 250 μl per well of DNA/RNA Shield buffer (Zymo R1100-250) was added. Plates were gently swirled until cells detached, and the solution pipette-mixed until homogeneous, then transferred to microcentrifuge tubes. RNA was extracted and concentrated using the RNA Clean & Concentrator (Zymo R1018) according to the manufacturer’s instructions. RNA was extracted similarly from 5-6 organoids per batch (n=8 differentiations from 6 cell lines) across DMSO and YTHDF inhibitor-treated groups (n=32 total samples; see “Cortical organoid culture and inhibitor treatment” below). For all KO and organoid samples, 500 ng total RNA was used to make RNA-seq libraries with the TruSeq Stranded Total RNA Library Prep kit (Illumina 20020599) following rRNA depletion (Illumina 20037135). Libraries were sequenced on an Illumina NovaSeq6000 to an average depth of 51M reads per library. Sequencing data were obtained for the following KO samples: FMR1 (n□=□8), FTO (n□=□5), METTL14 (n□=□8), METTL3 (n□=□7), WTAP (n□=□8), YTHDF1 (n□=□8), YTHDF2 (n□=□7), YTHDF3 (n□=□7), and NT (n□=□8).

### Bioinformatics analysis

To estimate the mutation rate introduced by the CRISPR system, we first aligned sequencing reads to the human reference genome (GRCh38, GENCODE Release 27) using STAR aligner^48^ (v2.6). Aligned reads were then categorized based on their CIGAR strings into two groups: perfect matches and reads containing mismatches, insertions, deletions, or clipping. The mutation rate was calculated by determining the proportion of reads with any mutations relative to the total number of aligned reads. Gene expression quantification was performed using Kallisto (v0.46) with annotations from GENCODE Release 27. Genes expressed in more than 50% of samples with at least 10 counts in each sample were retained for differential expression analysis, resulting in a total of 16,378 genes. Data normalization was carried out using the voom method from the limma package^51^, which models the mean-variance relationship in the log-counts. Differential expression between knockout (KO) and non-targeting (NT) guide RNAs (gRNAs) was tested using limma, controlling for sequencing batch, RNA preparation batch, and estimated library size.

### Western blot

At 9 dpt, media was aspirated from one well of a 6-well plate for each condition (8 KOs, 2 NT gRNAs, and mock-transfected/WT cells). Cells were washed once with 1X DPBS, then lysed in 250 μl in RIPA lysis buffer (Thermo 89900) containing cOmplete^TM^ Protease Inhibitor Cocktail (Roche 11697498001). Western blotting was performed using the Jess^TM^ Simple Western^TM^ (Biotechne 004-650) using the 12-230 kDa separation module (SM-W001). The Jess is an instrument enabling rapid/automated protein detection from low sample input. Lysate dilutions were optimized for each antibody and ranged from 1:3 to 1:9. Antibody dilutions were as follows: METTL3 1:20 (Abcam ab195352), METTL14 1:20 (Novus NBP1-81392), YTHDF2 1:20 (Abcam ab246514), YTHDF3 1:20 (Abcam ab220161), WTAP 1:20 (Abcam ab195380), FTO 1:40 (Abcam ab126605), Actin 1:50 (R&D MAB8929). Antibodies for YTHDF1 (ThermoFisher 17479-1-AP; Cell Signaling Technology 86463) and FMR1 (Biolegend 834701) were tested but did not produce a clean signal.

### Cortical organoid culture and inhibitor treatment

Human iPSC lines (BR62, BR63, BR01, BR21, BR22, BR28) were generated and characterized by the New York Stem Cell Foundation, and routinely maintained in mTeSR Plus medium (STEMCELL Technologies, #100-0276). The organoid differentiation was conducted based on published protocols with minor modifications^16^. In brief, iPSC were dissociated into single cells and aggregated at 10,000 cells/well of 96-well plate (S-Bio, MS-9096VZ) to form EBs in E6 medium (Fisher Scientific, A1516401) supplemented with Y-27632 (Tocris Bioscience, 1254) from day –1 to day 0. Neural induction (NIM phase) was initiated in E6 medium containing SB 431542 (10 μM) (Tocris Bioscience, #1614/10), LDN 193189 (100 nM) (Tocris Bioscience, #6053), and XAV 939 (2 μM) (Tocris Bioscience, #3748) from day 0 to day 4 to pattern EBs toward a cortical identity. From day 4 to day 10, XAV was withdrawn, while SB and LDN were continued. From day 10 to day 18, the medium was switched to neural differentiation medium (NDM phase) containing DMEM/F12, Neurobasal medium, N2 supplement, and B27 without vitamin A. From day 18 onward, 8 organoids were transferred to a single well of 6-well ultra-low attachment plate (Fisher Scientific, 07-200-601) on an orbital shaker and cultured in Neural Maturation Medium (NMM phase), supporting neuroepithelial rosette formation with twice-weekly media changes. The NMM medium consisted of DMEM/F12, Neural basal medium, N-2 supplement, B27 with vitamin A (Gibco, #17504-044), insulin (sigma, I9278), ascorbic acid (Cayman chemical, #16457), BDNF (ThermoFisher, #45002500UG), and dcAMP (Sigma, D0627).

Inhibitors of m6A readers were gifted from the laboratory of Dr Chuan He at the University of Chicago. The organoids (day 77) were treated with YTHDF1 inhibitor (SAC) at 20 μM, YTHDF2 inhibitor (DC-Y13-27) at 20 μM, combined SAC and DC-Y13-27 (both at 20 μM), and DMSO at 0.02% as the treatment control. 5 days later at day 82, organoids were collected for RNA extraction or fixed for 1 hour with 4% paraformaldehyde (Electron Microscopy Sciences, 15714-S) for immunohistochemistry.

For RNA-seq, organoids were generated from six iPSC lines derived from cognitively normal ROSMAP participants (BR62, BR63, BR01, BR21, BR22, BR28) across 1–2 independent differentiation batches, yielding *n* = 5–6 organoids per batch. For immunostaining, organoids were generated from two ROSMAP iPSC lines (BR62, BR63) across two independent differentiation batches, with 5–6 organoids per differentiation.

### Organoid IHC and image quantification

Fixed organoids were washed with 1x PBS, sunk in 30% Sucrose/1xPBS overnight, embedded in OCT, sectioned at 10 μm thickness, and mounted onto positive-charged glass slides, then stored at -80°C until staining. The sections were permeabilized in 0.3% Triton X-100 in 1xPBS for 30min, blocked in 5% BSA/1xPBS buffer for 1hr, and immunostained in primary antibodies diluted in blocking buffer at 4 °C overnight. Primary antibodies included PAX6 (DSHB, AB_528427), FOXG1(Abcam, ab18259), TBR2 (Abcam, ab23345), TBR1 (Abcam, ab31940), CTIP2 (Abcam, ab18465), SATB2 (Abcam, ab51502), MAP2 (Abcam, ab5392), and 4G8 (Biolegend, #800712). After washing three times with wash buffer (0.15% Tween-20/1xPBS), the antibodies were detected by Alexa Fluor-488 goat anti-chicken secondary antibody (ThermoFisher, A-11039) and Alexa Fluor-546 goat anti-mouse secondary antibody (ThermoFisher, A-11003) at 1:500 dilution in blocking buffer for 1 h at room temperature. After washing five times with the washing buffer, the sections were further counterstained with Hoechst nuclear stain (ThermoFisher, H21486) for 5 min, and coverslipped with ProLong™ Gold Antifade Mountant (ThermoFisher, P36934).

The images were acquired by ORCA-Flash4.0 V3 Digital CMOS camera in Nikon Eclipse Ti2 epifluorescence microscope with objective lens of 20x. Acquisition settings were the same for all the slides.

ND2 images were next imported to ImageJ for quantification. Regions with folds in the tissue or otherwise obvious technical artifacts were excluded for downstream analysis. For each image channel (DAPI, 4G8, and MAP2), we applied a minimum intensity threshold to exclude low/background signal, which was consistent across all images. We then analyzed particle metrics for each channel (e.g. area and puncta size), and generated summary metrics (total area, average puncta size, number of puncta). Summary metrics for 4G8 were normalized by MAP2 area to account for organoid size and neuronal composition. Data was imported to R for plotting and statistics. We used a nested ANOVA with post-hoc Tukey tests to determine whether groups exhibited significant differences.

### m6A eCLIP

m6A-eCLIP was performed at Eclipse BioInnovations Inc, San Diego CA as described below. The procedure started by cryogrinding approximately 200 mg of human brain tissue and isolating total RNA using a Maxi Prep kit (Qiagen). The RNA quality and quantity was assessed by Agilent TapeStation. rRNA was depleted by coupling 80 μg of Eclipse-designed probes with 600 μL of Streptavidin beads (ThermoFisher) for 10 minutes at room temperature. 10 μg of total RNA and pre-coupled bead-probes were incubated separately at 72°C for 30 seconds, then 67°C for 90 seconds. Without removing samples from incubator, the total RNA sample was combined with pre-coupled bead-probes and incubated at 64°C for 60 minutes. Following incubation, samples were magnetized, and the supernatant was reserved. RNA was cleaned using columns (Zymo) with a 200-nt size cut off. The mRNA quality and quantity was assessed by Agilent TapeStation. The mRNA was DNase-treated at 37°C for 10 minutes then immediately sheared into 100-200 nt fragments by heating at 95°C for 12 minutes in 1x Turbo DNase buffer (ThermoFisher). 4 μg of anti-m6A antibody (Eclipse Bio) was added, and samples were UV-C crosslinked using a UVP CL-1000 crosslinker at 2 rounds of 150 mJ/cm2 using 254 nm wavelength. The antibody-RNA complexes were then coupled overnight to protein G beads (CST). Following overnight coupling, library preparation (including adapter ligations, SDS-PAGE electrophoresis and nitrocellulose membrane transfer, reverse transcription, and PCR amplification) was performed as previously described for standard eCLIP^60^, with size-selection performed cutting the 30-110 kDa region from the membrane (corresponding to RNA fragments crosslinked to antibody heavy and light chains). 5% of fragmented mRNA was run as an RNA-seq input control, starting with FastAP treatment as described^60^. The final library’s shape and yield was assessed by Agilent TapeStation.

### Reader protein eCLIP

eCLIP studies were performed by Eclipse Bioinnovations Inc (San Diego, CA) according to the published single-end seCLIP protocol as described with the following modifications. Approximately 300 mg of frozen human brain tissue was cryogrinded and crosslinked using a UVP CL-1000 crosslinker at 2 rounds of 400 mJoules/cm2 using 254 nm wavelength. Cryogrinded tissue was separated into 30 mg aliquots for each protein type and then stored until use at -80°C. 30 mg of tissue was lysed using 1mL of eCLIP lysis mix with a modified composition of 6 μL of Proteinase Inhibitor Cocktail and 20 μL of Murine RNase Inhibitor. Samples were then subjected to 2 rounds of sonication for 4 minutes with 30 second ON/OFF at 75% amplitude. Eclipse validated antibodies YTHDF1(ProteinTech: Cat No. 17479-1-AP), YTHDF2 (Aviva Systems Biology: Cat No. ARP67917_PO50), YTHDF3 (Abcam: Cat No. ab220161), and FMRP (Bethyl: Cat No. A305-200A) were then pre-coupled to Anti-Rabbit IgG Dynabeads (ThermoFisher 11203D), added to lysate, and incubated overnight at 4°C. Prior to immunoprecipitation, 2% of the sample was taken as the paired input sample, with the remainder magnetically separated and washed with eCLIP high stringency wash buffers. Following overnight coupling, library preparation (including adapter ligations, SDS-PAGE electrophoresis and nitrocellulose membrane transfer, reverse transcription, and PCR amplification) was performed as previously described for standard eCLIP^60^ with size-selection performed cutting the region from 65 kDa to 140 kDa for YTHDF1, 2 and 3 and from 65 kDa to 150 kDa for FMRP. RNA was visualized using Chemiluminescent Nucleic Acid Detection Module kit (ThermoFisher 89880).

### eCLIP analysis

The eCLIP cDNA adapter contains a sequence of 10 random nucleotides at the 5’ end. This random sequence serves as a unique molecular identifier (UMI)^61^ after sequencing primers are ligated to the 3’ end of cDNA molecules. Therefore, eCLIP reads begin with the UMI and, in the first step of analysis, UMIs were pruned from read sequences using umi_tools (v1.1.0)^62^. UMI sequences were saved by incorporating them into the read names in the FASTQ files to be utilized in subsequent analysis steps. Next, 3’-adapters were trimmed from reads using cutadapt (v3.2)^63^, and reads shorter than 18 bp in length were removed. Reads were mapped to the human genome (hg38) using STAR (v2.6.1d)^48^. PCR duplicates were removed using umi_tools (v1.1.0) by utilizing UMI sequences from the read names and mapping positions. Reads mapping to exons of GENCODE (Release 27) and not mapping to human repetitive elements (identified using the RepeatMasker track in the UCSC Genome Browser) were retained for further analysis. Peaks were identified within eCLIP samples using the peak caller CLIPper (v2.0.0) ^64^. To reduce the computational time required to apply CLIPper to all samples, analysis was restricted to genes expressed in the input samples across m6A, YTHDF1, YTHDF2, YTHDF3, and FMR1 eCLIP experiments. Gene expression was quantified using featureCounts (v2.0.1)^65^, and 11,633 genes with at least 10 counts in 50% of the samples were retained for CLIPper peak detection. For each peak, the enrichment of IP versus input was calculated using Fisher’s Exact Test based on a 2×2 contingency table comparing read counts within the peak region between IP and input samples. To control for multiple testing, Storey’s q-value method^66^ was applied, and peaks with an FDR ≤5% were retained for each sample. Consensus peaks were defined as regions detected in at least 40 samples. These regions were then extended to include adjacent regions supported by at least 20 samples, and overlapping regions were merged to form the final consensus peak set. Read coverage for these consensus peaks was calculated across all samples using featureCounts (v2.0.1) with default parameters. For finer m6A site resolution, single nucleotide sites were also called. In this analysis, 5′ mismatches were trimmed from read alignments to remove potential artifacts. Single-nucleotide resolution crosslink sites were then called from the trimmed reads using PureCLIP v1.3.1^67^, utilizing input control data to identify sites enriched in the IP over the input. Enrichment of IP versus input at these sites was calculated using Fisher’s Exact Test as described above, and the q-value method^66^ was applied to control for multiple testing, retaining sites with an FDR ≤□5% in each sample. Consensus sites were defined as sites detected in at least 40 samples. Read counts overlapping these consensus sites were then quantified using bedtools (v2.30.0)^68^.

### Peak locations along regional annotations

We visualized peak locations relative to transcript models using the Guitar package (v2.4.0)^69^. To ensure computational efficiency and maintain clarity in visualization, we randomly selected up to 10,000 peaks for analysis when the total number of peaks exceeded this threshold. Peak annotations were assigned based on their genomic regions, including promoters, exons, introns, and intergenic regions, to contextualize the binding sites within the transcript architecture.

### Motif analysis

We conducted de novo motif identification using HOMER^70^ (v4.11) to search for enriched sequence motifs of length 5 base pairs (bp) within consensus peaks identified by CLIPper for each RNA-binding protein (RBP): YTHDF1, YTHDF2, YTHDF3, FMR1, and m6A eCLIP datasets. Given that de novo motif identification is a stochastic process and may yield variable results across different datasets, we aggregated all identified de novo motifs to create a comprehensive set for further analysis. To evaluate the enrichment of these aggregated motifs across all experimental conditions, we utilized the monaLisa package^71^ (v1.0.0) in R. We selected the top two motifs from each eCLIP dataset based on statistical significance and assessed their enrichment across all antibodies using consistent parameters. To ensure a fair comparison between different datasets, we standardized the peak regions by extracting sequences of fixed length—100 bp centered around the peak summits—for all analyses.

### Differential peak analysis

Normalization of IP read counts for consensus peaks was conducted using DESeq2^72^ (v1.34.0), which models count data based on the negative binomial distribution. We adjusted for RNA Integrity Number (RIN), post-mortem interval (PMI), and sequencing batch effects in both IP read counts and gene counts in input samples. To account for IP efficiency and parental RNA expression levels, we followed the procedure outlined in a previous study^73^. Specifically, peaks within the top 1% of counts ranked by IP read counts were normalized by their corresponding parental gene counts. Correction factors for IP efficiency differences were then estimated by computing the median of each column after normalizing each row by the geometric mean. These correction factors were applied to the IP read counts to normalize for IP efficiency. To further account for differences in parental gene expression across samples, we calculated the ratio of gene expression differences against the mean expression across all samples for each gene. IP read counts were adjusted by dividing by this ratio for each gene, thereby normalizing for variations in parent gene expression levels. Differential peak analysis between AD and NCI samples was performed using DESeq2, with adjustments for sex, age at death, and years of education as covariates. False discovery rate (FDR) was estimated using Storey’s q-value method. Peaks were considered significantly differentially expressed if they met thresholds of p < 0.05 and FDR < 0.05.

### Predicting AD diagnosis from eCLIP and RNA-seq data

We performed elastic net regression using the glmnet package in R (version 4.1) to model the relationship between AD diagnosis and eCLIP as well as RNA-seq data from the DLPFC. Peak intensities from IP eCLIP data were normalized by the corresponding parental RNA expression levels to account for variability in RNA abundance. To reduce the number of variables in the elastic net model, we first identified co-expressed modules using the SpeakEasy algorithm with default settings (v1.0.0)^57^. For RNA expression data, we utilized the module definitions from our previous study^1^. Gene expressions were standardized by subtracting the mean and dividing by the standard deviation for each gene across all samples. An average module expression was then computed by taking the mean of the standardized expressions of all genes within each module. Model performance was assessed using a 5-fold cross-validation approach. Performance metrics included root mean square error (RMSE) and Pearson’s correlation coefficient between predicted and observed values. Hyperparameter tuning was conducted for the mixing parameter alpha, ranging from 0.1 to 1.0 in increments of 0.1, and the regularization parameter lambda, tested over a logarithmic scale from 1×10^−6^ to 100. The optimal model was selected based on the combination of alpha and lambda that minimized the RMSE and maximized the correlation between predicted and observed values. The model outputs the probability of a sample being AD-positive. We converted these probabilities to log odds and computed the area under the receiver operating characteristic curve (AUC-ROC) and the area under the precision-recall curve (AUC-PR) to evaluate model performance. All performance evaluations were conducted using test samples that were excluded from any model training processes. The PRROC package (v1.3.1) and the pROC package (v1.18.5) in R were used to compute these metrics.

## Materials availability

This study did not generate new unique reagents.

## Data availability

All data is available upon request through the RADC Resource Sharing Hub (www.radc.rush.edu) and the AMP-AD Knowledge Portal in accordance with the data usage agreement. The ROSMAP RNA-seq data from the dorsolateral prefrontal cortex (DLPFC) is available for download from the AMP-AD Knowledge Portal under Synapse accession ID syn3388564. The ROSMAP TMT proteomics data from frozen DLPFC tissue can be accessed from the Synapse repository under Synapse accession IDs syn30390636, syn32539359, and syn59611693. eCLIP data is available for download from the AMP-AD Knowledge Portal under Synapse accession ID syn68174819. Gene expression data from CRISPR knockout experiments will be available on the Gene Expression Omnibus (GEO) under accession ID GSE286096. The AD Knowledge Portal is a platform for accessing data, analyses, and tools generated by the Accelerating Medicines Partnership (AMP-AD) Target Discovery Program and other National Institute on Aging (NIA)-supported programs to enable open-science practices and accelerate translational learning. The data, analyses, and tools are shared early in the research cycle without a publication embargo on secondary use. Data is available for general research use according to the following requirements for data access and data attribution (https://adknowledgeportal.org/DataAccess/Instructions). RNA-binding regions are available at POSTAR2 (http://postar.ncrnalab.org). Transcription factor binding regions are available at GTRD (https://gtrd.biouml.org).

## Code availability

The DEcode-tree and eCLIP-seq analysis pipelines used in this study will be publicly available on GitHub(https://github.com/stasaki/DEcodeTree and https://github.com/stasaki/eCLIPseq-pipeline).

## Acknowledgments

We are grateful to those who agreed to donate their brains for research. We thank all the employees at the Rush Alzheimer’s Disease Center (RADC) for their support and assistance. This study was supported by faculty start-up funding (to Y.W.) and NIA grants R01AG074082 and R01AG079223 (to Y.W.), P30AG10161, R01AG015819, R01AG017917, and U01AG061356 (to D. A. B.), R01AG061798 and R01AG057911 (to C.G.), R01AG072120 (to A.P.W and T.S.W), R01AG022018 (to L.L.B.), R01AG095602 (C.H. and Y.W.), P30AG072975, R01AG057911, U01AG079847, R01AG15819 (ROSMAP; genomics and RNAseq), R01AG30146, R01AG36836 (RNAseq), R01AG034374, R01AG042210, R01AG072120, RF1AG036042, RF1AG015819, RF1AG074549, U01AG32984 (genomic and whole exome sequencing), U01AG046152, U01AG46161, U01AG061357 (TMT proteomics), and P30AG010161. Data collection was also supported by the Illinois Department of Public Health and the Translational Genomics Research Institute. These data were provided through the Accelerating Medicine Partnership for AD (U01AG046161) based on samples from the Rush Alzheimer’s Disease Center. Data were also provided by Dr. Levey from Emory University, based on postmortem brain tissue collected through the Mount Sinai VA Medical Center Brain Bank provided by Dr. Eric Schadt from Mount Sinai School of Medicine. The results published here are in whole or in part based on data obtained from the AD Knowledge Portal (https://adknowledgeportal.synapse.org). Study data were provided by the Rush Alzheimer’s Disease Center, Rush University Medical Center, Chicago. Additional phenotypic data can be requested at www.radc.rush.edu.

## Author contributions

S.T. performed all computational analysis. D.R.A. performed gene perturbation and RNAseq experiments and analyzed organoid staining. N.K. generated Cas9-expressing cell lines. A.I., H.V., C.Y. and D.J.F. performed pilot immunohistochemistry experiments to validate the predictions. W.L. and C.H. graciously provided YTHDF inhibitors. C.Y. and L.C. performed organoid culture and treatment with inhibitors. S.D.T and J.X. helped with sequencing experiments. S.T., D.R.A., and Y.W. wrote the manuscript with input from all co-authors. L.L.B runs the parent Minority Aging Research Study and African American Clinical Core. D.A.B runs the parent ROSMAP study. S.T. and Y.W. conceived and supervised this study. All authors reviewed and approved the final manuscript.

## Declaration of interests

G.W.Y. is an SAB member of Jumpcode Genomics and a co-founder, member of the Board of Directors, on the SAB, equity holder, and paid consultant for Eclipse BioInnovations. G.W.Y.’s interests have been reviewed and approved by the University of California, San Diego, in accordance with its conflict-of-interest policies.

---

**Supplementary Fig. 1.**
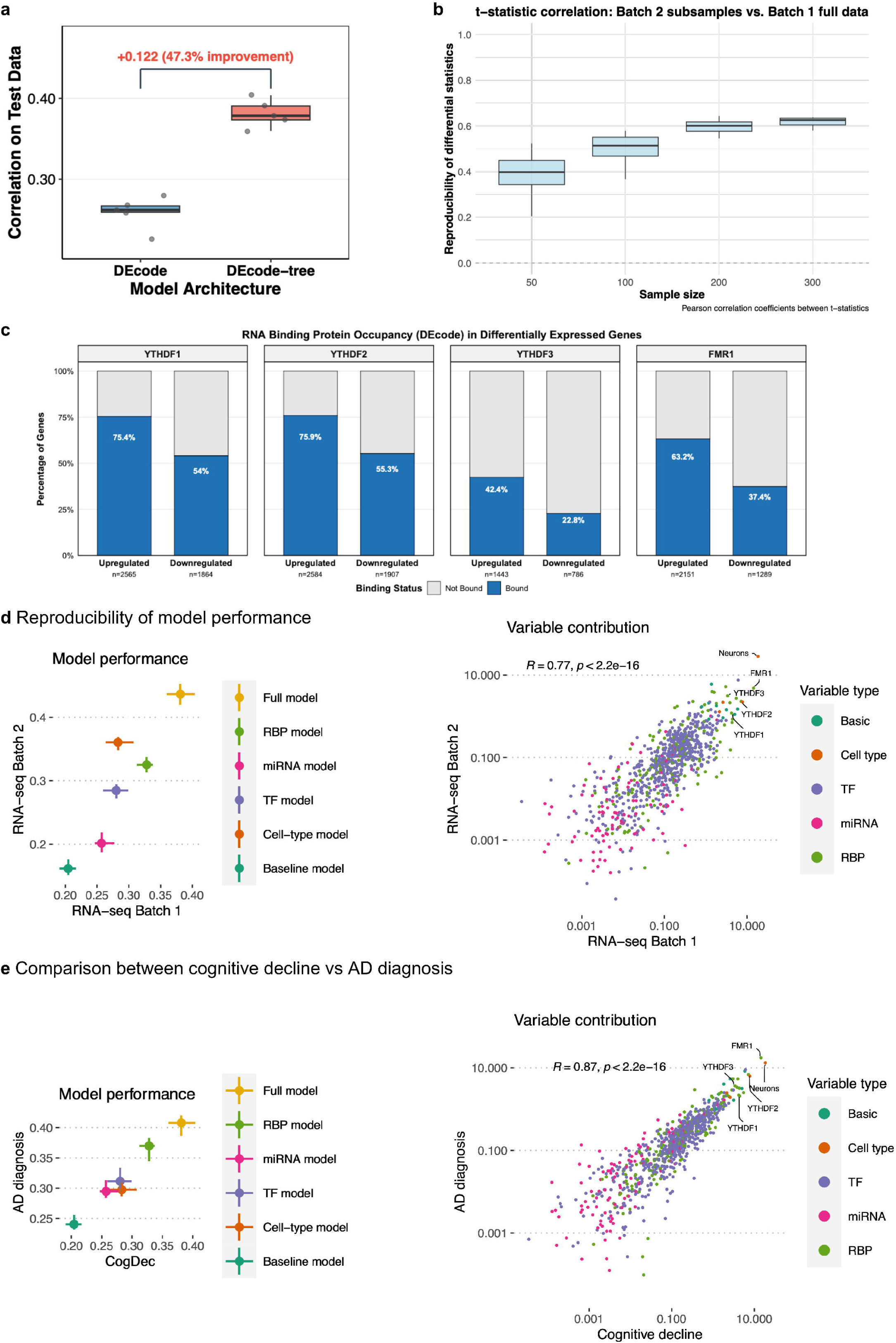

**Supplementary Fig. 2.**
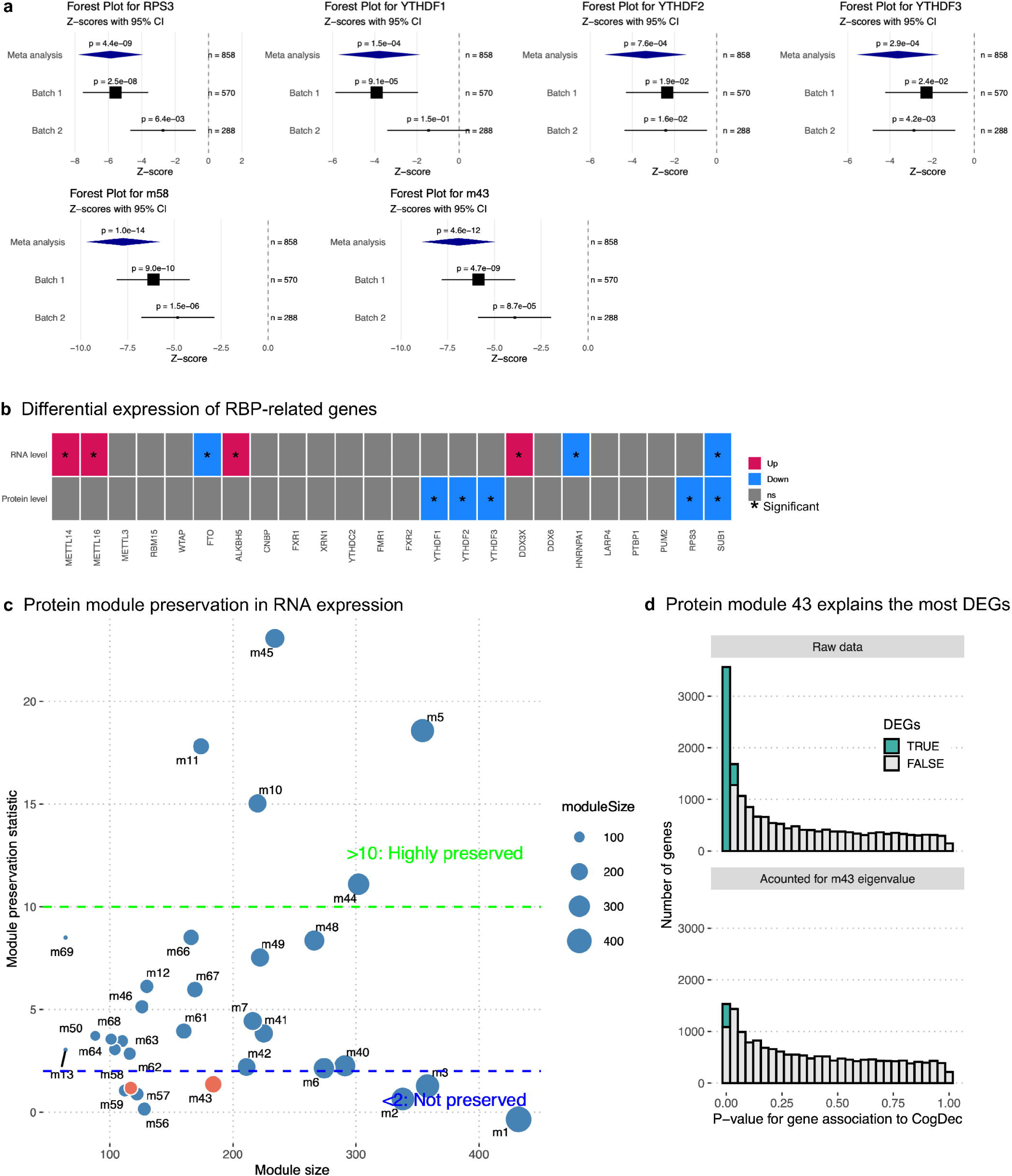

**Supplementary Fig. 3.**
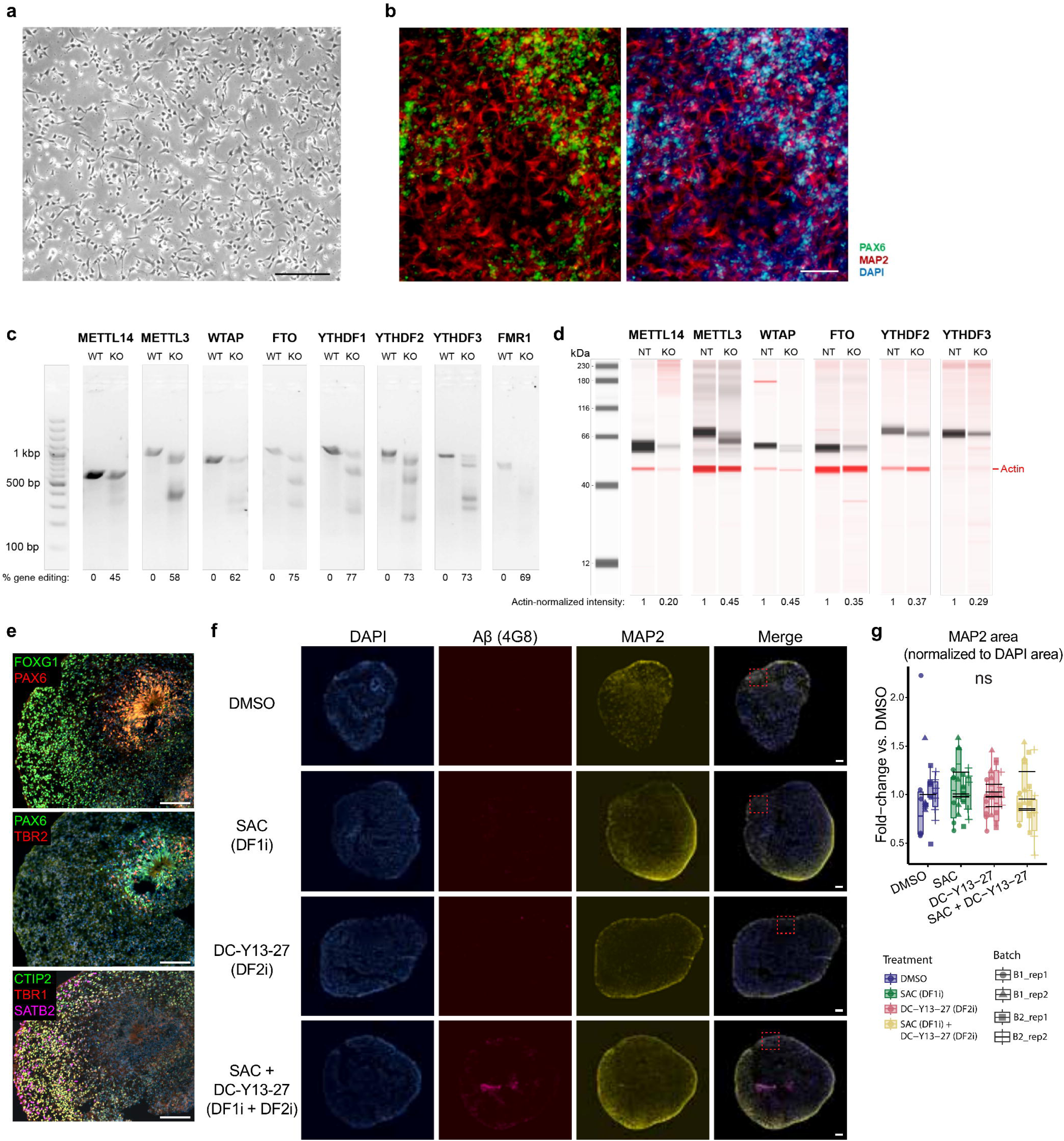

**Supplementary Fig. 4.**
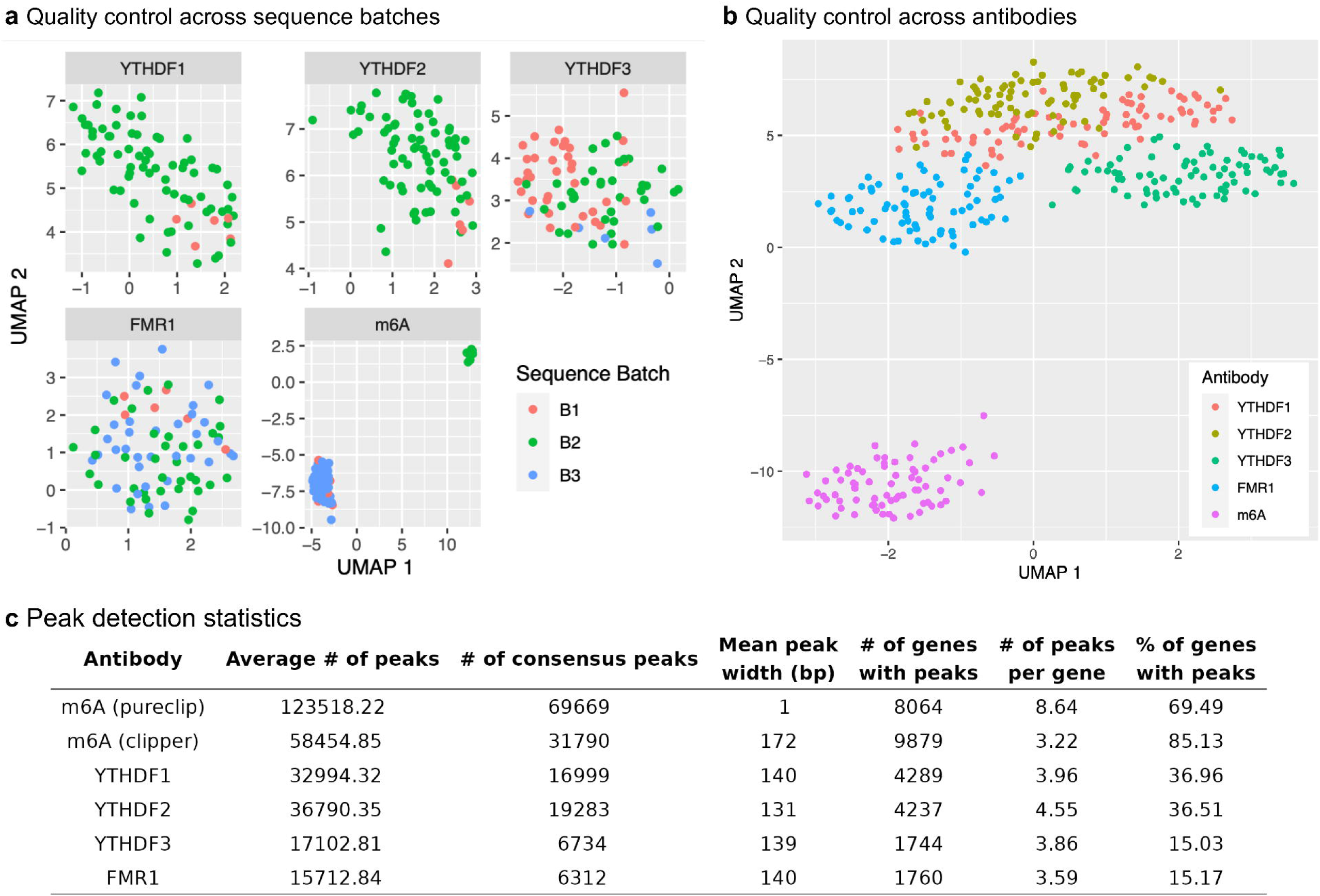

**Supplementary Fig. 5.**
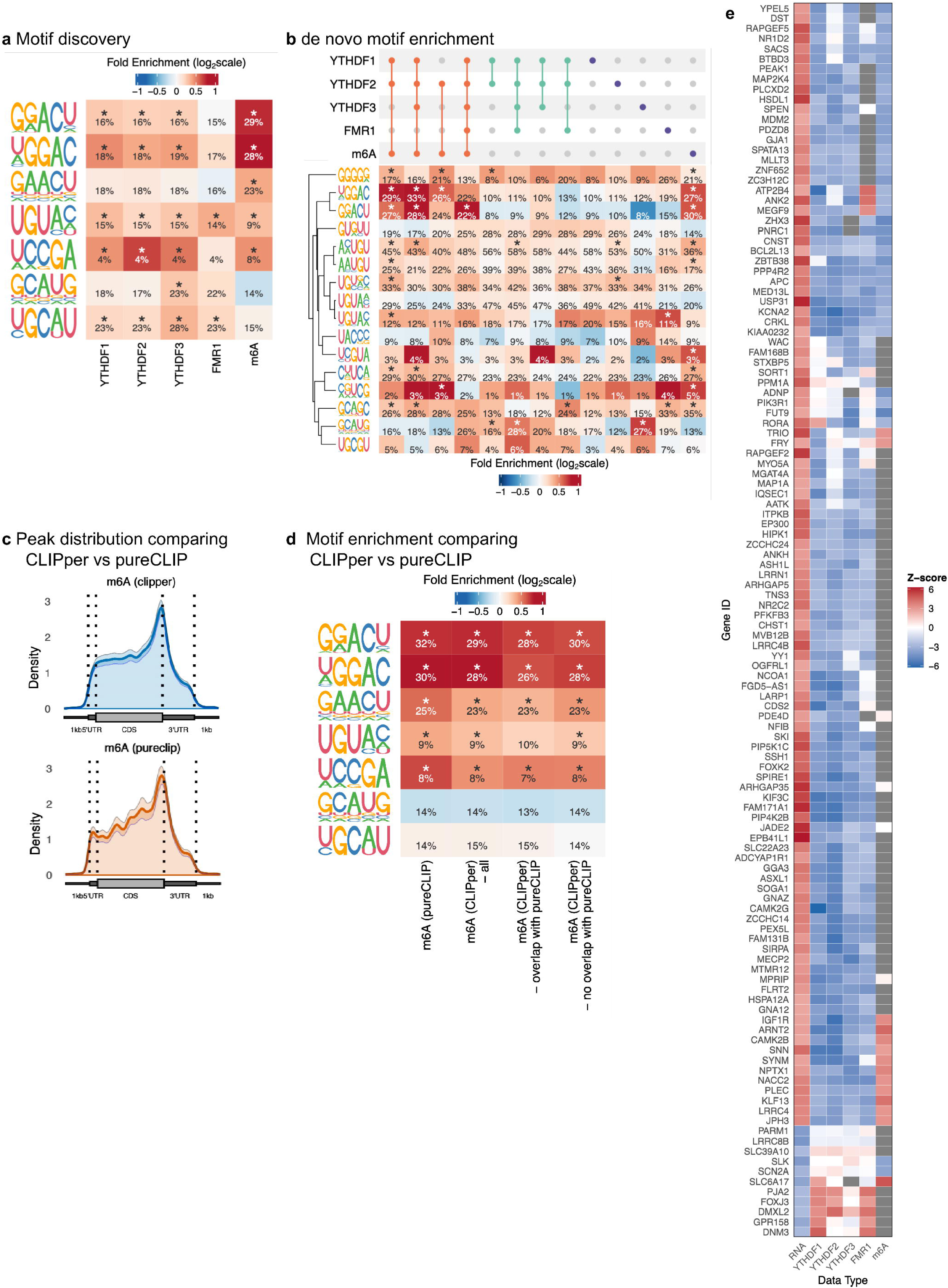

## Notes

### Competing Interest Statement

The authors have declared no competing interest.

### Summary of Updates

This version of the manuscript has been revised to include additional analyses (for example, using a new batch of proteomics data), new pharmacological inhibition experiments to further validate the findings, and an updated author list reflecting changes in authorship.

